# Calcineurin controls the cytokinesis machinery during thermal stress in *Cryptococcus deneoformans*

**DOI:** 10.1101/2025.02.17.638697

**Authors:** Vikas Yadav, Anna Floyd Averette, Rajendra Upadhya, Joseph Heitman

## Abstract

Calcineurin is a highly conserved phosphatase that plays a central role in sensing calcium and governing transcriptional, post-transcriptional, and post-translational signaling networks. Calcineurin is a heterodimer consisting of a catalytic A subunit and a regulatory B subunit. Through downstream effectors, calcineurin signaling drives myriad responses in different organisms. In the fungal pathogenic *Cryptococcus* species complex that infects humans, calcineurin governs thermotolerance and is essential for growth at high temperature and pathogenesis. In *Cryptococcus deneoformans*, the underlying molecular functions of this critical signaling cascade are not well understood. In this study, we conducted a genetic screen and identified genetic changes that suppress the requirement for calcineurin during high-temperature growth. Our results identified two mechanisms that bypass the requirement for calcineurin function. The first mechanism involves segmental aneuploidy via both amplification as well as loss of chromosome fragments. The second mechanism involves dominant amino acid substitution mutations in the genes encoding three proteins, Chs6, Imp2, and Cts1, orthologs of components of the Ingression Progression Complex required for septation and budding in *Saccharomyces cerevisiae*. Loss of calcineurin activity causes chitin and chitosan accumulation and severe budding defects, whereas suppressor mutations largely restore growth and cytokinesis in the absence of calcineurin. These findings reveal the calcineurin signaling cascade controls a conserved cytokinesis machinery at the mitotic exit network during thermal stress.

**Significance Statement:** Cellular responses to external stress conditions involve activation of signaling cascades, which act via effectors to govern key cellular processes. One such signaling cascade involves the phosphatase calcineurin, the conserved target of cyclosporin A and FK506 that governs myriad cellular functions through its substrates. In human fungal pathogens, calcineurin is critical for pathogenicity. Here, we employed a genetic screen to map the functional interactions of this signaling cascade and repeatedly identified mutations in components of a single protein complex, suggesting a focused downstream response. With genetic and biochemical approaches, we established that these proteins play a crucial role in cell division at high temperature enabling cell division and revealing new avenues of research that could identify potential antifungal drug targets.

## Introduction

Stress adaptation is crucial for all organisms to grow and survive in adverse conditions. Unicellular organisms exposed to such conditions employ a diverse range of strategies to adapt to stress. Stress adaptation strategies are critical for pathogenic organisms to survive host body temperature as well as to tolerate chemical and nutrient stress conditions. Thus, the ability to grow at human body temperature (37°C) is essential for any systemic human pathogen. Despite this, the molecular mechanisms underlying thermal adaptation are incompletely understood in most human pathogens, including eukaryotic pathogens such as fungi. As climate change challenges our planet, understanding the ways in which microbes, including pathogenic ones, respond to and adapt to thermal stress will be paramount in controlling the threats that they present.

The *Cryptococcus* species complex consists of a group of related human fungal pathogens that account for >110,000 deaths each year and ∼20% of HIV/AIDS-related deaths worldwide (1). While *Cryptococcus neoformans* is the primary global pathogen in this species complex, *Cryptococcus deneoformans* and *Cryptococcus gattii* infections are geographically restricted and more prevalent in certain regions of the world (2, 3). Thermotolerance is an established virulence attribute for these pathogens and mutants exhibiting defects in thermotolerance are attenuated or avirulent. Previous studies have established that calcium-calcineurin signaling is crucial for thermotolerance in this ubiquitous fungal pathogen (4, 5).

Calcium-calcineurin signaling is a highly conserved cascade among eukaryotes that is activated upon calcium influx into the cytoplasm (6–9). Calcineurin is a heterodimeric phosphatase comprising two subunits, the catalytic subunit A (Cna1) and the regulatory subunit B (Cnb1). Upon influx into the cytoplasm, calcium binds to calmodulin (CaM) as well as to the regulatory subunit of calcineurin. This is followed by the association of CaM with the calcineurin heterodimer, which leads to conformational changes, release of the autoinhibitory domain from the active site, and activation of the phosphatase. Calcineurin then acts via downstream effector proteins, including conserved transcription factors (NFAT in humans and Crz1 in fungi) (10–13). Through these effectors, calcineurin plays a crucial role in the immune system in humans and stress responses in fungi. Calcineurin activity can be inhibited by FK506 or cyclosporin A (CsA), both of which are gold standard FDA-approved immunosuppressive drugs in widespread clinical use (11, 14). In *C. neoformans*, calcineurin effectors were previously identified to implicate a major role for calcineurin in thermal adaptation through functions executed at processing bodies (P-bodies) and stress granules (10, 15–17). A potential role for calcineurin at the mother-bud neck has also been implicated in this species (15). In contrast to *C. neoformans*, the underlying mechanisms of calcineurin function in governing thermotolerance remain unknown in *C. deneoformans* and other related species (5).

In this study, two genetic screens were conducted to define calcineurin functions in *C. deneoformans*. First, spontaneous mutations that restore growth at 37°C in the presence of calcineurin inhibitors, FK506 and/or CsA, were isolated. Second, spontaneous mutations were isolated that restore growth at 37°C of a mutant lacking calcineurin regulatory subunit. Both of these approaches resulted in the isolation of more than 50 independent suppressors that bypass the requirement for calcineurin during growth at 37°C. Whole genome sequencing followed by variant calling identified mutations in several genes, of which three were identified in independent isolates. Additionally, segmental aneuploidy was also found to bypass the requirement for calcineurin for 37°C growth. Further analysis revealed that dominant mutations in proteins involved in cytokinesis bypass calcineurin with high efficiency providing evidence that calcineurin plays a major role during cell division in this species. Overall, our results provide insights into calcineurin’s role in cell division and suggest the presence of a mitotic exit network that is conserved with *Saccharomyces cerevisiae*.

## Results

### Mutations that bypass calcineurin function in C. deneoformans

To understand the basis of calcineurin function, we employed two different approaches to isolate spontaneous genetic mutants. In the first approach, we plated the wild-type congenic strains JEC20**a** and JEC21α on media containing both calcineurin inhibitors, FK506 and CsA, and incubated at 37°C to obtain suppressor colonies in which high temperature growth is no longer inhibited by FK506 and CsA (Fig. 1A). We also isolated some mutants on only FK506-containing media that were then found to be cross-resistant to CsA at 37°C, and these were included in the analysis. Notably, double mutations in both FKBP12 and cyclophilin A (required for FK506 or CsA toxicity) are expected to be exceedingly rare. Moreover, mutations in calcineurin A or B that prevent FKBP12-FK506 binding have not been found to cause CsA resistance (18). In the second approach, we utilized calcineurin mutant strains, JEC20**a** *cnb1*Δ and JEC21α *cnb1*Δ, and isolated colonies growing at the restrictive temperature of 37°C. Several independent plating experiments were conducted for each strain and condition and 64 independent suppressor mutants were identified. We confirmed the ability of these mutations to suppress calcineurin function by growing them under restrictive growth conditions, in the presence of calcineurin inhibitors at 37°C for the wild-type strains and at 37°C for the calcineurin mutant strains (Fig. 1B). All of the suppressor mutant strains were able to grow under restrictive conditions whereas the parental strains were not.

**Fig. 1.**
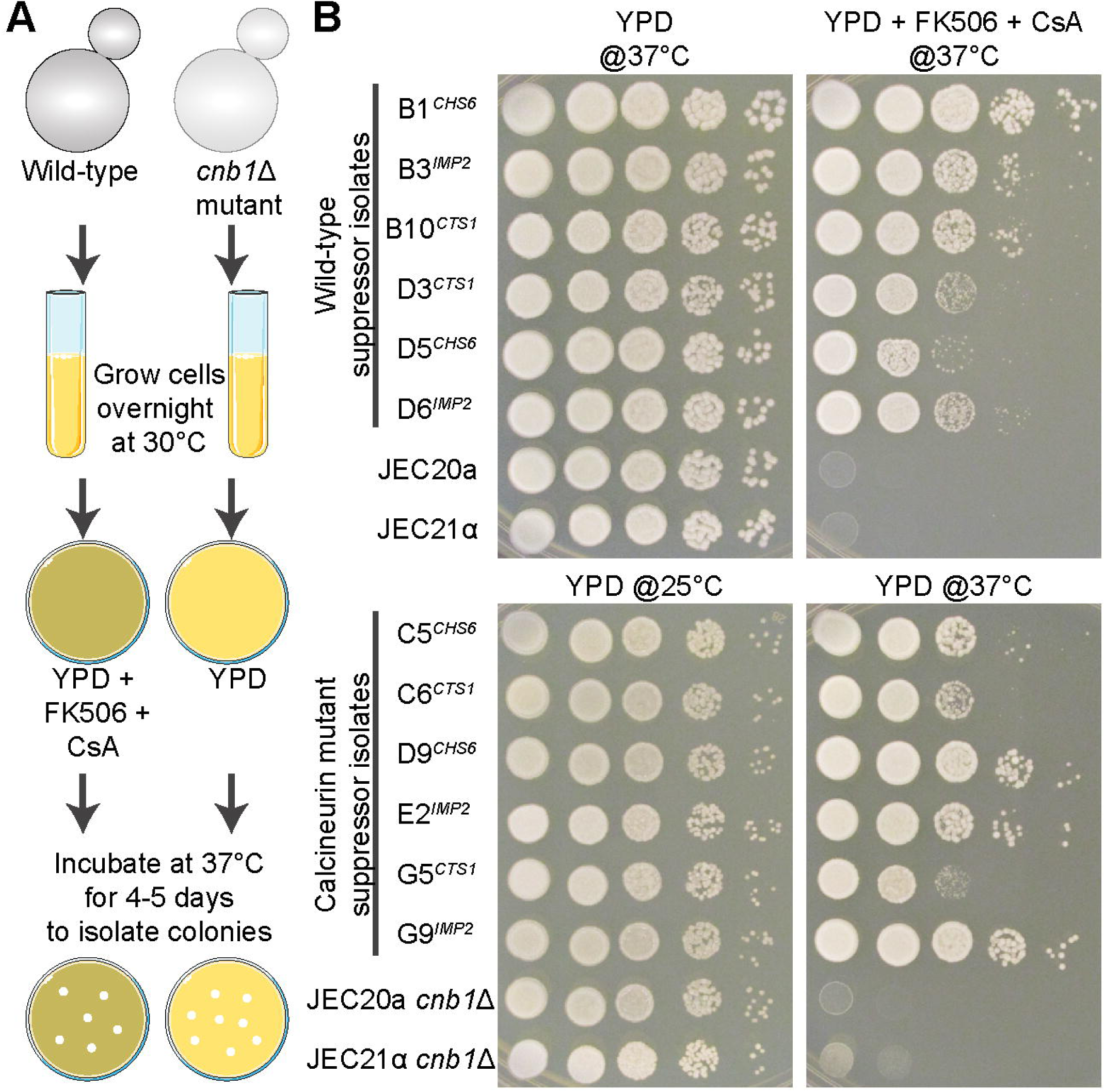
Identification of calcineurin suppressors in *C. deneoformans*. **(A)** A scheme depicting the approach employed to isolate calcineurin suppressor mutants in both the wild-type and calcineurin deletion mutant strains. **(B)** Plate images showing the growth of calcineurin suppressor isolates in restrictive conditions compared to the parental strains. YPD+FK506+CsA at 37°C is the non-permissive condition for the wild-type JEC20 and JEC21 strains, whereas YPD at 37°C is the non-permissive condition for the calcineurin mutants, JEC20 *cnb1*Δ and JEC21 *cnb1*Δ. CsA is cyclosporin A; Gene names are listed as superscripts next to the isolates listed (i.e. B1, B3).

In addition to high temperature growth, calcineurin is also necessary for sexual reproduction in *C. deneoformans* (4, 5, 19). Calcineurin mutants are bilaterally sterile, meaning that both mating partners must lack calcineurin for a defect in sexual hyphae to be observed. We tested whether these suppressors were able to also bypass calcineurin function in sexual reproduction by crossing both the wild type and the suppressor strains to the respective opposite mating type partners. The wild-type strains as well as suppressor mutants formed robust hyphae when crossed with partners on Murashige and Skoog (MS) mating media indicating that the suppressor mutations had no impact on mating (SI Appendix, Fig. S1). However, none of the suppressors examined showed mating hyphae in the presence of FK506, similar to the wild type. Because several of these suppressors are dominant (shown below), in these crosses the wild type partner lacks calcineurin (due to FK506 inhibition) but the *cna1* mutant partner with the dominant suppressor (not complemented by cell fusion) has calcineurin’s role at high temperature bypassed, if a common target controlled both high temperature growth and sexual development, mating should have been restored. Because mating was not restored, we conclude that the isolated mutations specifically suppress calcineurin’s role in thermal stress adaptation but not in sexual reproduction. Thus, independent calcineurin regulatory pathways govern thermal tolerance and mating.

Next, these isolates were subjected to Illumina whole genome sequencing to identify the underlying genetic mutations. Sequence data were processed and mapped to the wild-type reference genome, JEC21α, and variant calling was performed to identify causative genetic changes. Surprisingly, a large number of suppressor isolates harbored only one genetic change in a single protein-coding region for each isolate, suggesting the presence of one determinant factor in each strain. In other isolates, no clear genetic changes in protein-coding regions were identified. A subset of these strains was subjected to further analysis and scored for mutations in intronic regions that could potentially affect gene function as well as aneuploidy. By these approaches, we were able to assign a single causative genetic change in each of the suppressors that ranged from amino acid substitution mutations to large-scale aneuploidy (Table 1 and Dataset S1). Several of these aneuploid chromosomes and mutations were shared between wild-type suppressor isolates cross-resistant to FK506+CsA and *cnb1*Δ mutant suppressors suggesting the presence of a mechanism shared between these two groups. Overall, this analysis revealed that in *C. deneoformans* mutations in several independent genes can suppress the normal requirement for calcineurin to grow at high temperature.

**Table 1.**
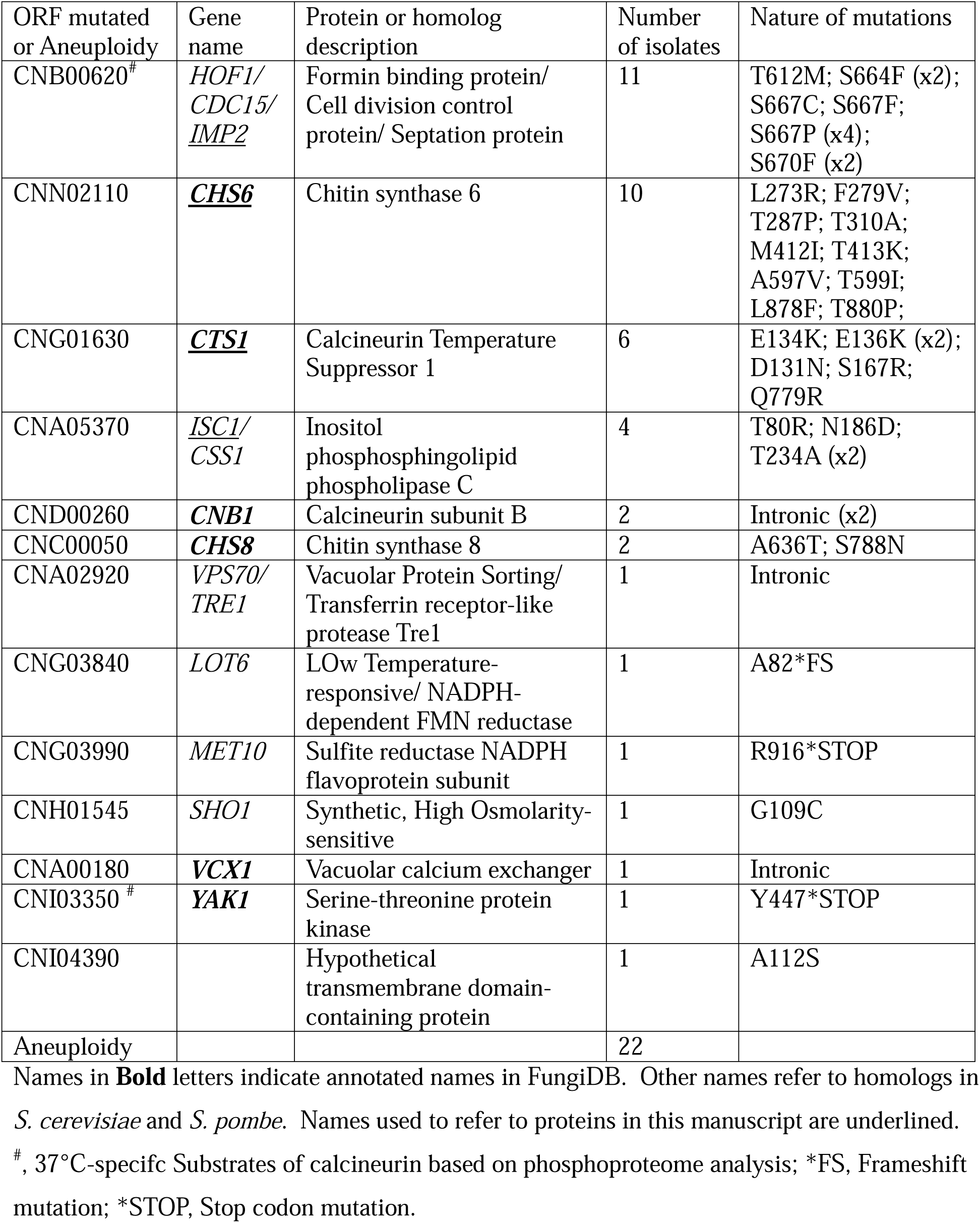
Genes mutated in calcineurin suppressor isolates.

### Segmental aneuploidy plays a major role in calcineurin functional suppression

Among the calcineurin suppressor mutations, segmental aneuploidy was the largest determinant factor with 22 out of 64 strains harboring either an amplification or a loss of part of a chromosome. Interestingly, several independent regions were found to be aneuploid with four of these shared among at least three independent isolates (Fig. 2A). Analysis of the underlying genes in these four regions identified candidate genes in two regions based on their described roles in thermotolerance and calcineurin signaling. For chromosome 7, the aneuploid region harbors a gene that was previously identified as a multicopy suppressor of calcineurin function for high temperature growth in *C. deneoformans* and named Calcineurin Temperature Suppressor 1 or *CTS1* (Fig. 2A, middle panel) (20). The Cts1 protein contains a C2 domain, which is required for high temperature growth, virulence, and hyphal growth, and Cts1 functions in a parallel pathway with calcineurin. In *C. neoformans*, Cts1 localizes with calcineurin at the mother-bud neck and P-bodies and is also a substrate of calcineurin (21). Combined with these previous studies, segmental amplification of *CTS1* leading to overexpression is the most likely cause of calcineurin function suppression in these isolates.

**Fig. 2.**
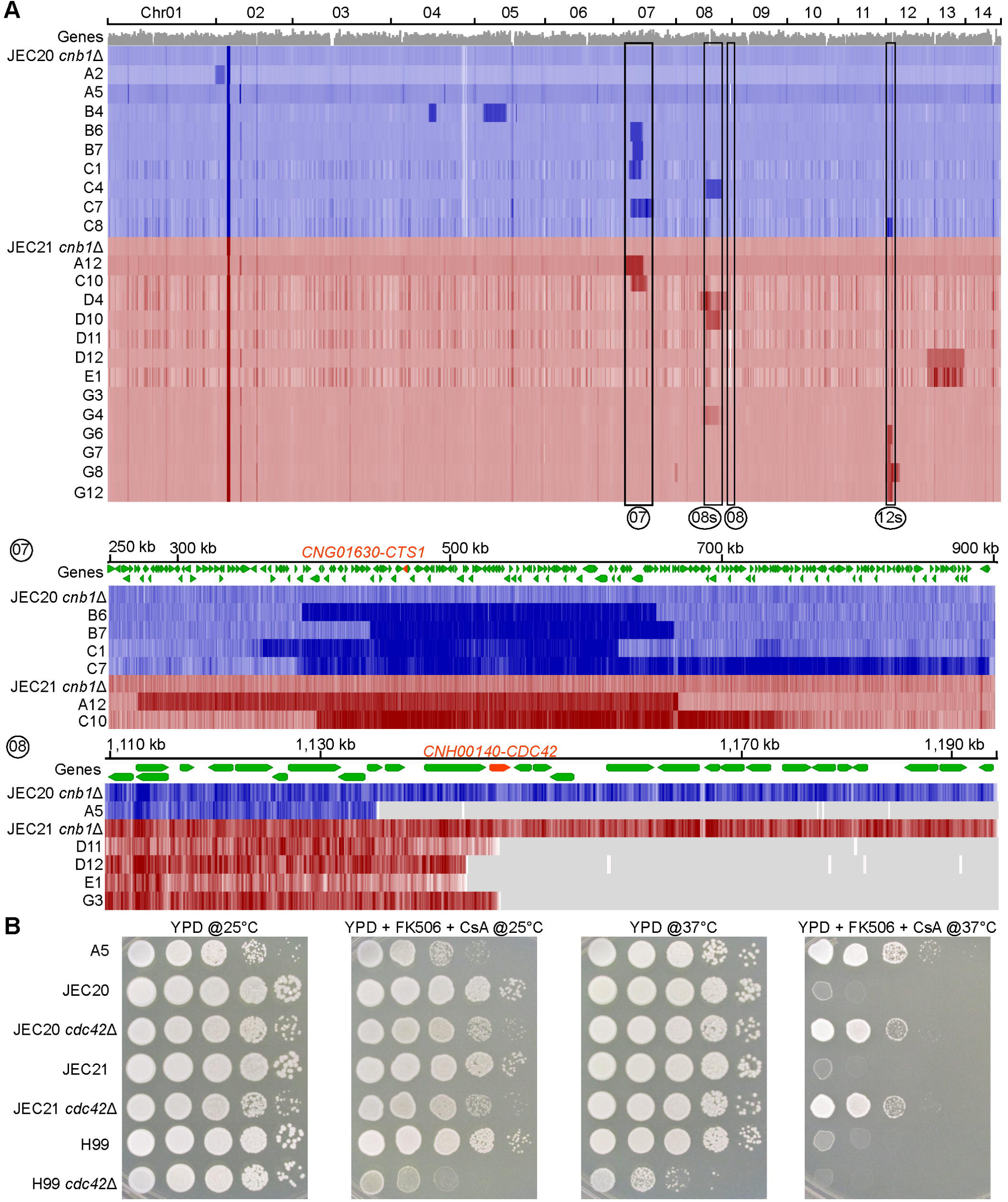
Segmental aneuploidy suppresses loss of calcineurin function and restores high temperature growth. **(A)** A heatmap showing coverage of the reference JEC21 genome from the whole genome sequencing for selected isolates (22 isolates with aneuploidy and 2 controls). All strains derived from JEC20 are presented in blue whereas JEC21-derived strains are in red. Each row represents sequencing data from one strain, JEC20 *cnb1*Δ and JEC21 *cnb1*Δ exhibited uniform coverage throughout the genome as controls. The remaining rows present suppressor isolates with at least one event of aneuploidy marked by either darker color (amplification) or loss of colored regions (loss of a fragment) in heatmaps. The middle and lower panels show the zoomed view of two such regions marked as 07 (for chromosome 07) and 08 (for chromosome 08) in the upper panel and only show isolates with the mentioned aneuploidy. *CTS1* (*CNG01630*) and *CDC42* (*CNH00140*) are highlighted as the potential candidate genes that could drive calcineurin bypass in regions 07 and 08, respectively. The regions marked as 08s and 12s in the upper panel are presented in SI Appendix, Fig. S1. A complete list of genes present in each aneuploid region is presented in the Dataset S2. **(B)** Plate images showing serial dilution spotting of wild-type JEC21 and JEC20 strains with their respective *cdc42*Δ mutants along with the suppressor A5 that exhibited aneuploidy causing loss of Cdc42. *C. neoformans* wild-type H99 and its respective *cdc42*Δ mutants were generated in a previous study (22) and are shown for comparison.

For chromosome 8, we observed two separate aneuploidy events among different isolates. One of these caused the loss of one end of the chromosome with *CDC42*, a gene encoding for a small rho-like GTPase, either entirely lost or partially truncated (Fig. 2A, lowest panel). Cdc42 has been previously shown to play an important role in budding and cell division in *C. neoformans* and is also required for thermotolerance (22). Interestingly, our results indicate that loss of Cdc42 confers an ability to grow at 37°C in the case of *C. deneoformans* in the absence of calcineurin, in contrast to *C. neoformans* where *cdc42*Δ mutant itself is temperature sensitive. To validate that loss of Cdc42 is indeed the causative factor for calcineurin suppression at high temperature, we genetically deleted *CDC42* in *C. deneoformans* strains JEC20 and JEC21. Deletion of Cdc42 led to growth restoration in the presence of calcineurin inhibitors to the same extent as that of the aneuploid isolate A5 (Fig. 2B). These results provide strong evidence that the loss of Cdc42 is responsible for bypassing calcineurin function in these aneuploids. Notably, both aneuploid A5 and *cdc42* deletion mutants exhibited slower growth in other tested conditions, suggesting a fitness defect. We also tested *cdc42*Δ mutants in *C. neoformans*, which grew poorly at 37°C and did not suppress calcineurin inhibition. Interestingly, loss of Cdc42 in *C. neoformans* also made cells hypersensitive to calcineurin inhibitors even at 25°C suggesting that calcineurin and Cdc42 function in parallel pathways. These findings also suggest a rewiring of functional links between Cdc42 and calcineurin has occurred between these two species. Both Cts1 and Cdc42 play a role in cell budding and septation indicating that calcineurin plays an important role in cell division and either overexpression of Cts1 or deletion of Cdc42 could enable cell division to be completed in the absence of calcineurin.

For the other two genomic regions that were aneuploid in multiple isolates as well as other aneuploid regions that were present in one or two isolates, the causative genes were unclear as these regions contained several genes encoding hypothetical proteins and proteins involved in various cellular processes (Dataset S2, SI Appendix, Fig. S2). Segmental aneuploidy has been shown to provide resistance to several drugs in *Cryptococcus* species, including azoles (23, 24). We identified segmental aneuploidy that could provide resistance to the calcineurin inhibitors, FK506 and CsA. However, the same regions of aneuploidy were also identified in suppressors where calcineurin B was genetically deleted, indicating that aneuploidy plays a broader role in providing adaptation to stressful conditions by enabling bypass of the requirement for calcineurin for high temperature growth.

### Mutations in IMP2, CHS6*, and* CTS1 are dominant

Apart from the aneuploid events, three genes harbored multiple independent mutations in more than five isolates (Fig. 3A). The gene that harbored the greatest number of mutations, 11 in total, encodes an F-BAR domain-containing protein with an SH3 domain at the C-terminus and a highly disordered central region. The encoded protein is an ortholog of Hof1 in *S. cerevisiae* and Imp2/Cdc15 in *Schizosaccharomyces pombe* and is referred to as Imp2 in this study (25–28). In both *S. cerevisiae* and *S. pombe*, Imp2 orthologs play a crucial role in septation during cell division (25, 29). Our screen identified several independent mutations in this protein, all of which are serine/threonine mutations suggesting that these mutations could impact the phosphorylation status of the protein.

**Fig. 3.**
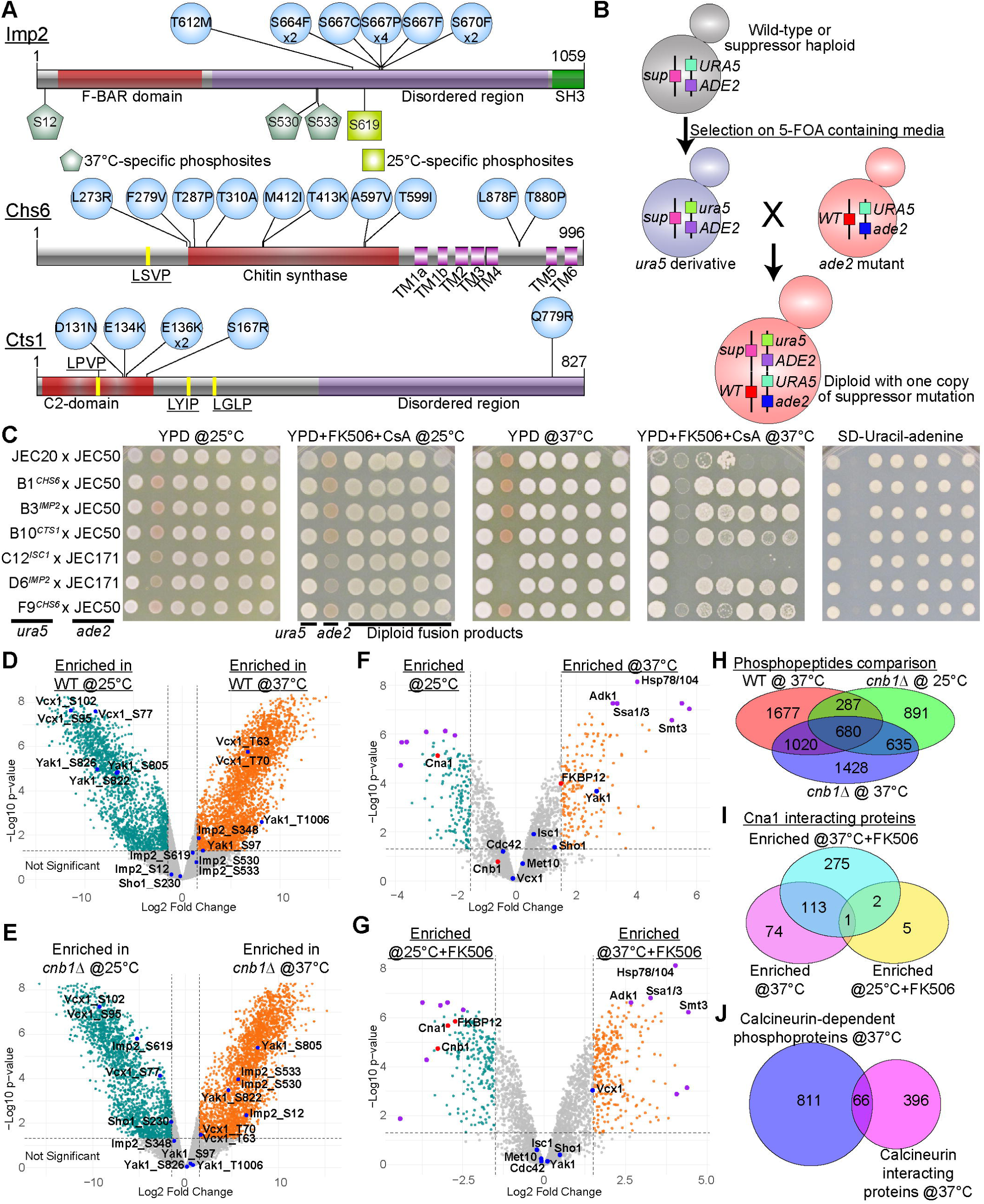
Mutations in Imp2, Chs6, and Cts1 are dominant. **(A)** Cartoon representations of three proteins that acquired multiple independent mutations are shown with their domain architecture. All identified mutations are amino acid substitutions and are marked as circles on their respective positions. LSVP on Chs6 and LPVP, LYIP, and LGLP on Cts1 were identified as variants of the LxVP motif as potential docking motifs for calcineurin. TM1a-TM6 in Chs6 refer to the transmembrane domains identified in this protein. Pentagon-shaped sites on Imp2 are 37°C-specific calcineurin-mediated phosphosites whereas rectangles mark 25°C-specific phosphosites. **(B)** A scheme showing the outline to generate diploid strains from identified suppressors to determine the dominant-recessive phenotype of the mutations. **(C)** Plate images showing the growth of generated diploids from the wild-type strain JEC20 as well as various suppressors in the presence of calcineurin inhibitors FK506 and cyclosporin A (CsA) at 37°C revealed the dominant nature of mutations identified in *CHS6*, *IMP2,* and *CTS1*. Growth on standard-defined (SD) media lacking uracil and adenine for diploids is shown as a control for the presence of both functional *URA5* and *ADE2* genes in the heterozygous diploids. **(D-E)** Phosphoproteome analysis identified phosphosites that were enriched in the wild-type and calcineurin mutants at 25°C and 37°C and pair-wise comparison is presented as volcano plots. Significantly enriched peptides (log2 fold change >1.5; p-value < 0.05) are colored in each case. Phosphosites in proteins that bypassed calcineurin and were significantly enriched in at least one group are highlighted in volcano plots. **(F-G)** Volcano plots presenting the proteins identified from the Cna1-GFP immuno-pulldown assays at 25°C and 37°C with or without FK506. Pair-wise comparisons are shown with the identified suppressors positions marked in blue and expected proteins (Cna1, Cnb1, and FKBP12) in red, and the 6 most significantly enriched proteins in purple. Names for some of the most significantly interacting proteins that were shared at 37°C between with or without FK506 conditions are also shown. **(H)** A Venn diagram presenting the overlap among phosphosites that were enriched in three groups (wild type at 37°C, *cnb1*Δ at 25°C and *cnb1*Δ at 37°C) as compared to wild type at 25°C. **(I)** A Venn diagram showing the overlaps of significantly Cna1-GFP interacting proteins as compared to the 25°C without the FK506 control group. **(J)** A cartoon showing the overlap of proteins corresponding to the 37°C-specific calcineurin-dependent phosphopeptides (877 proteins corresponding to 1428 phosphopeptides shown in H) and proteins that were found to interact with Cna1-GFP only at 37°C (combined pool from with or without FK506 conditions).

The next gene with 10 different mutations encodes a chitin synthase, Chs6, which belongs to the chitin synthase class I/II (30). The majority of the mutations were confined to its glycotransferase or chitin synthase domain indicating that these mutations likely impact its activity. Notably, we also identified mutations in *CHS8*, encoding another member of the class I/II chitin synthase family indicating that mutations in either of the two members can lead to calcineurin suppression in this species. Interestingly, class I/II chitin synthases are also known to play a role in cell division in *S. cerevisiae* as well as in *Candida albicans* (31, 32).

The third gene with 6 mutations encodes Cts1, which is the same gene that was also present in one of the aneuploid groups described earlier. 5 of these mutations are present in the C2 domain and may, therefore, impact its function. As noted, Cts1 was shown to localize to the mother-bud neck in *C. neoformans* and plays a role in budding (20, 21).

All of the mutations identified in *CHS6*, *CTS1,* and *IMP2* cause amino acid substitutions (Table 1). Combined with calcineurin bypass via *CTS1* copy number increase through aneuploidy, these results indicate a potential gain of function for the mutated proteins. Furthermore, suppressor mutations in all of these genes were identified not only in the wild-type background but also in the calcineurin mutants suggesting that these are bypass mutations for calcineurin’s function and do not require calcineurin to be present for their action. These lines of evidence suggest that these mutations may be dominant. To test this, we generated diploids that harbored one copy of the suppressor mutation and one copy of the wild-type gene (Fig. 3B). First, we obtained uracil auxotrophic mutants for the calcineurin suppressors as well as for the wild-type JEC20 by selecting on 5-Fluoroorotic acid (5-FOA) media. Next, we crossed these to adenine auxotrophic strains of the opposite mating type and selected prototrophic fusion diploids on media lacking uracil and adenine. The resulting diploids were confirmed by flow cytometry and the presence of both mating type alleles by PCR analysis. The confirmed diploids were tested for ability to grow in the presence of calcineurin inhibitors FK506 and CsA at 37°C. Interestingly, diploids carrying the *CHS6*, *IMP2*, and *CTS1* mutations exhibited robust growth under this otherwise restrictive condition, even in the presence of a wild-type allele of the gene (Fig. 3C and SI Appendix, Table S1). The diploids with two wild-type alleles did not exhibit any growth on media containing FK506 and CsA at 37°C, as expected. These results demonstrate that the *CHS6*, *CTS1,* and *IMP2* suppressor mutations are dominant and thus may provide a gain of function. We also tested this phenotype for another suppressor gene that encodes a phospholipase C (*ISC1*) and found this specific mutation is recessive as diploids carrying the mutation and a wild-type gene copy failed to grow at the restrictive condition.

### Imp2 is a substrate of calcineurin

Chs6 and Cts1 both harbor one or more LxVP calcineurin docking motifs indicating that these proteins may be substrates of calcineurin (Fig. 3A) (33, 34). Indeed, Cts1 was established as a substrate of calcineurin in *C. neoformans* in a previous study (21). Interestingly, several of the suppressor mutations impact serine or threonine residues that might be dephosphorylated by calcineurin in the wildtype proteins (5/10 mutations in Chs6, 1/6 in Cts1, and 11/11 in Imp2). To determine whether one or more of these proteins could be direct substrates of calcineurin, phosphoproteome analysis was conducted with whole-cell lysates for both the wild-type and *cnb1*Δ mutant cells at 25°C and 37°C. PCA analysis and heatmap clustering analysis revealed that three replicates in each condition were strongly correlated (SI Appendix, Fig. S3A-B). The phosphopeptides that were enriched in the calcineurin mutant specifically at 37°C were identified and analyzed (Fig. 3D-E, 3H, SI Appendix, Fig. S3C and Dataset S3). This resulted in the identification of 1428 phosphopeptides corresponding to 877 proteins. Gene ontology analysis of these 877 proteins revealed a role of calcineurin in several biological processes and molecular functions (SI Appendix, Fig. S3C-D). Interestingly, cellular component-based analysis of 877 proteins revealed a significant enrichment of proteins that are present in the cell cortex and cell tip suggesting a role for calcineurin in cell polarity and growth, which might be relevant to its role during sexual reproduction (SI Appendix, Fig. S3D-F).

Among these proteins, neither Chs6 nor Cts1 were identified as substrates in this analysis despite having one or more LxVP motifs. However, Imp2 was identified as a candidate substrate for calcineurin with at least 3 phosphosites that were unique to 37°C in the *cnb1*Δ mutant (S12, S530, and S533) and one phosphosite that was unique to 25°C (S619) (Fig. 3A and D-E). In addition, one more 37°C-specific calcineurin-independent phosphosite was also identified (S348). Importantly, these sites differ from those whose mutation resulted in calcineurin bypass. Based on this data, two hypotheses might explain the results. First, Chs6, Cts1, and Imp2 might exist as a complex in the cell and either Chs6 or Cts1 might recruit calcineurin through their LxVP motifs, which then leads to dephosphorylation of Imp2. Second, the phosphoregulation of these proteins by calcineurin may occur at different times and our analysis may not be under the optimal conditions to identify temporally distinct events. It is possible that the whole proteome level phosphoproteome was unable to identify all of the phosphosites in Imp2 as well as any modification in Chs6 and Cts1.

In addition to Imp2, the phosphoproteome analysis also identified 37°C-specific calcineurin-dependent dephosphorylation of Yak1, mutations of which are also associated with calcineurin suppression. Interestingly, the mutation in Yak1 is likely a loss of function because it leads to an in-frame stop codon mutation. We hypothesize that the Yak1 kinase might be responsible for the phosphorylation of certain proteins that are dephosphorylated by calcineurin for high-temperature growth. Thus, loss of Yak1 could restore the proteins to an unphosphorylated state, mimicking activated calcineurin and thus allowing the growth of cells.

### Calcineurin localizes at the mother bud neck

To further elucidate the mechanism of calcineurin function suppression through mutations in Chs6, Cts1, and Imp2, we tested whether these proteins directly interact with calcineurin, possibly at the mother-bud neck. For this purpose, Cna1 was tagged at the C-terminus with GFP and co-localized with Cdc10-mCherry. Cdc10 is a septin that localizes to the mother bud neck in *C. neoformans* (35) and a similar localization pattern was observed in *C. deneoformans* in our study. Consistent with the previously shown localization of Cna1 in *C. neoformans* at the mother bud neck (21), Cna1 localized at the mother bud neck partially overlapping with the Cdc10 signal during cell division at 37°C and exhibited a dynamic localization in *C. deneoformans* as well (SI Appendix, Fig. S4). We also attempted to tag and localize Chs6, Cts1, and Imp2 but were unable to localize GFP-tagged versions of Cts1, Imp2, or Chs6 successfully.

We then conducted immunoprecipitation assays of the Cna1-GFP fusion protein with GFP-TRAP beads both in the presence and absence of FK506 to determine whether Cna1 physically associates with Cts1, Imp2, and/or Chs6. The immunoprecipitation assays were done with cells grown at 25°C and 37°C to identify thermal stress-specific interactions. All replicates for each condition showed strong clustering in PCA and heatmap analyses (SI Appendix, Fig. S5), and expected proteins (Cna1, Cnb1, and FKBP12) were significantly enriched in all datasets. Interestingly, a larger number of interacting proteins were identified in the presence of FK506 at 37°C than without FK506 (Fig. 3F, 3G, and 3I). However, the two sets exhibited significant overlap suggesting that the presence of FK506 might stabilize calcineurin interactions. We also identified potential direct substrates of calcineurin by combining Cna1-interacting proteins with phosphoproteome data (Fig. 3J). This analysis resulted in the identification of 66 proteins that directly interact with calcineurin and are also dephosphorylated in a calcineurin-dependent manner suggesting that these are direct calcineurin substrates (SI Appendix, Table S2). Interestingly, only Yak1 was found to be a direct interacting protein among the suppressors and none of the Chs6, Cts1, or Imp2 were detected in the mass-spec data (Fig. 3F). A lack of these proteins from immunoprecipitation data suggests that the interaction may be transient or indirect among these proteins.

Gene ontology (GO) analysis of these 66 proteins did not reveal enrichment of any specific biological process or cellular component. However, several proteins involved in mRNA binding/processing, mitochondria-related functions, and multiple heat shock proteins (Hsp) were identified in this pool. Previously, calcineurin was shown to localize to P-bodies and act on several mRNA-binding proteins at these subcellular organelles in *C. neoformans* (15, 17). Interestingly, Cts1 was also shown to localize to P-bodies in *C. neoformans* and it is possible that Cna1 directly interacts with Cts1 at these structures, even though our data failed to identify this specific interaction (21). Our data revealed calcineurin interactions with Hsp70 and its cochaperones suggesting the formation of calcineurin-Hsp70 complex that could govern its function. Calcineurin has been shown to interact with Hsp90 in *C. albicans*, specifically as a client of Hsp90 (36) whereas its interactions with Hsp70 are not well-defined. Hsp70 is involved in the folding and homeostasis of several proteins, and it may regulate several proteins involved in budding and cell polarity including Chs6, Cts1, and Imp2.

### CHS6, CTS1, and IMP2 suppressor mutations allow cytokinesis in the absence of calcineurin activity

Homologs of Chs6 and Imp2 play important roles during cytokinesis in *S. cerevisiae* and *S. pombe*. Additionally, Cts1 in *C. neoformans* also localizes to the mother-bud neck. This prompted us to hypothesize that all of these proteins are involved in cell division, specifically in cytokinesis. The processes of cell division and septation have been well studied in *S. cerevisiae* and, interestingly, a key set of proteins required for proper cytokinesis includes an F-BAR domain-containing protein Hof1 (ortholog of Imp2), a C2-domain containing protein Inn1 (ortholog of Cts1), and a chitin synthase Chs2 belonging to the class I/II family (ortholog of Chs6) (26, 37–40). These three proteins along with Cyk3 form a complex, known as the Ingression Progression Complex (IPC), which is involved in the formation of primary septa during cytokinesis (41, 42). Furthermore, these proteins physically interact with each other in a way that results in the activation of Chs2, which is crucial for chitin synthesis during primary septa formation (26, 28, 38, 43).

To determine whether Chs6, Cts1, and Imp2 interact with each other, we predicted their structures using AlphaFold 3 (Fig. 4A and SI Appendix, Fig. S6) (44). AlphaFold modeling of the Chs6 homodimer revealed a structure very closely resembling the cryoEM structures of *C. albicans* Chs2 (45) and *S. cerevisiae* Chs1(46). Mapping of the identified mutations on the structure revealed that all of the mutations were localized either in the domain swap region or the adjacent glycosyltransferase domain in the neighboring protomer (SI Appendix, Fig. S6A-B). Interestingly, this domain swap region has been proposed to play a key role in the activation of chitin synthesis activity. Modeling of Chs6, Cts1, and Imp2 together showed the formation of a hexamer complex (2 copies of each protein) with an interface of the C2-domain of Cts1 with the N-terminal of Chs6 (Fig. 4A and SI Appendix, Fig. S6). Furthermore, Chs6, Cts1, and Imp2 all exhibit a high level of disorder in their predicted structures when analyzed on their own, but the disordered regions were structured in the multimer predicted complex suggesting that interaction of these proteins with each other could stabilize the complex and promote folding of individual subunits (SI Appendix, Fig. S6A-E). This analysis suggests that a similar machinery may also be present in *C. deneoformans*, a basidiomycetous yeast, and our result identifying calcineurin-dependent Imp2 phosphorylation sites implicates regulation by calcineurin for its proper function at 37°C.

**Fig. 4.**
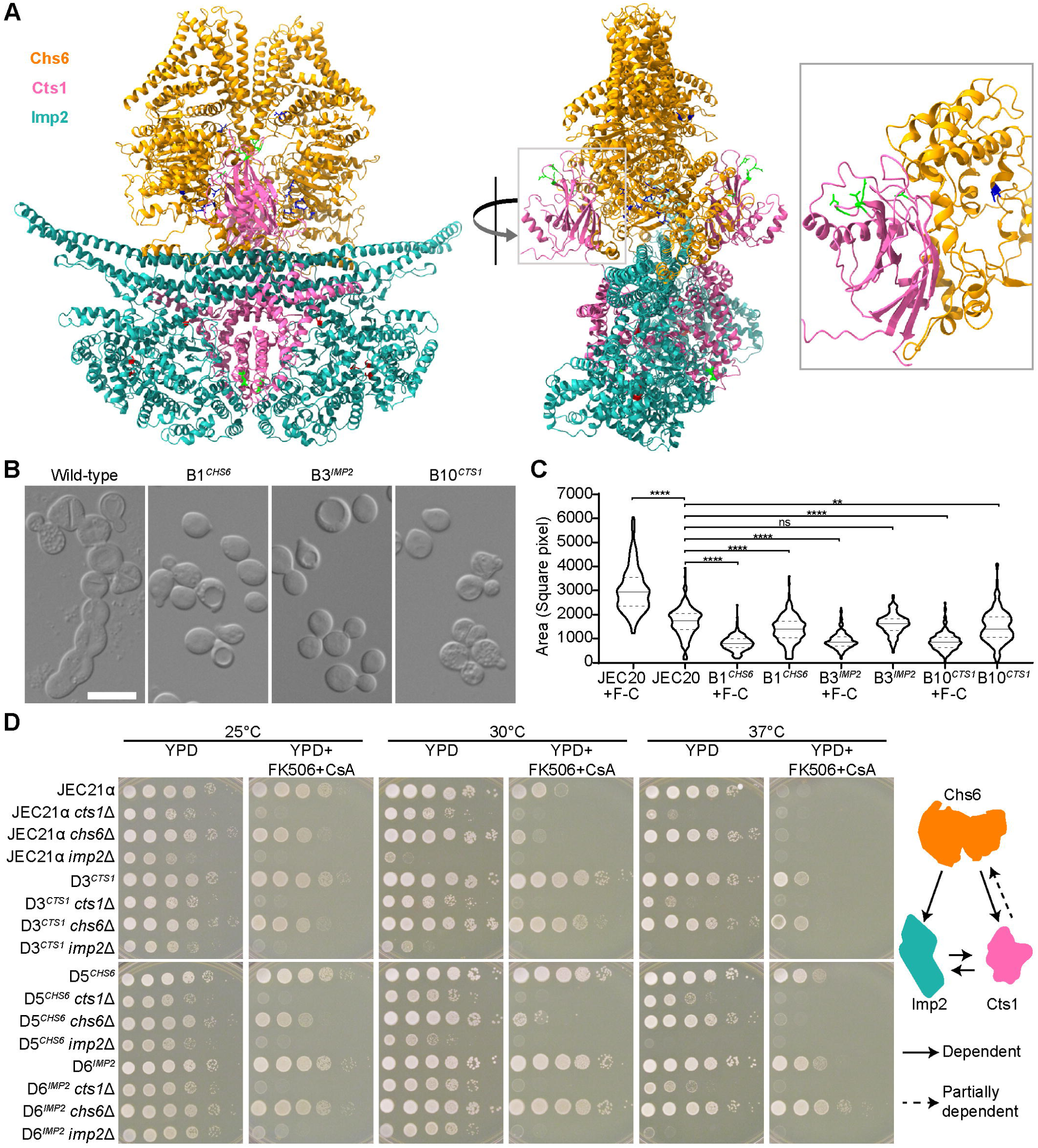
Imp2, Chs6, and Cts1 form an inter-dependent complex that drives cytokinesis. **(A)** An AlphaFold 3 structure prediction of the Chs6-Cts1-Imp2 complex in a dimer configuration revealed a strong complex formation between these three proteins. Identified calcineurin suppressor mutations are marked in blue for Chs6, green for Cts1, and red for Imp2. The inset panel shows a zoomed view of the interaction between the C2-domain of Cts1 (hot pink) with the structured N-terminal region of Chs6 (orange), which is highly disordered otherwise (SI Appendix, Fig. S5). **(B)** DIC images showing the cell morphology of wild-type and strains with suppressor mutations in Chs6, Cts1, and Imp2 after growth in media containing FK506 and CsA. Scale bar, 5 µm. **(C)** Cell size was quantified in wild-type JEC20 and its bypass mutants (B1. B3 and B10) in both treated and untreated conditions at 37°C. Cell size was measured in brightfield images as the surface area of cells (n > 150 cells) and is presented as square pixels. The solid horizontal line marks the median whereas dotted lines represent the first and third quartiles. F-C refers to samples treated with FK506 and cyclosporin A. **, p<0.01; ****, p<0.0001; ns, non-significant. **(D)** Serial dilution spotting assays showing the growth of *chs6*Δ, *cts1*Δ, and *imp2*Δ mutants in the wild-type JEC21 and suppressor strains with mutations in Chs6, Cts1, or Imp2. Based on their growth levels in respective deletion mutants, Chs6, Cts1, and Imp2 were found to be interdependent, with the exception that Imp2 was not dependent on Chs6, which is shown by the cartoon representation.

To study this, we determined the cellular morphology of wild-type and suppressor mutants after a prolonged inhibition of calcineurin activity. The untreated wild-type cells showed normal morphology with a mixed population of unbudded and budding cells (SI Appendix, Fig. S7A) whereas cells treated with both FK506 and CsA exhibited severe budding defects exhibiting long chains of cells that failed to undergo cytokinesis (Fig. 4B and SI Appendix, Fig. S7A). Chitin staining in these cells revealed a uniform staining along the cell wall as well as a bright septation site in untreated budding cells (SI Appendix, Fig. S7A). In contrast, the FK506 + CsA treated cells not only exhibited a very strong signal at the mother bud neck, but this signal was quite elongated consistent with a broadened neck region in these cells. This was further confirmed with scanning electron microscopy (SEM) that clearly showed a cytokinesis defect with a broadened neck region between the undivided cells (SI Appendix, Fig. S8). In addition, these cells showed patches of chitin accumulation in other parts of the cell wall, suggesting dysregulated chitin synthesis. Quantification of total chitin and chitosan levels revealed the wild-type strain JEC20 accumulated higher levels of chitin when treated with calcineurin inhibitors FK506 and CsA (SI Appendix, Fig. S7B).

The *CHS6*, *CTS1,* and *IMP2* suppressor mutations largely restored cellular budding and chitin levels despite the presence of calcineurin inhibitors (Fig. 4B and SI Appendix, Fig. S7A). SEM imaging further revealed successful budding events in cells with a *CHS6* suppressor (B1) mutation even in the presence of calcineurin inhibitors (SI Appendix, Fig. S8). Measurement of chitin and chitosan levels in suppressor mutants showed similar levels of these polymers in both treated and untreated conditions supporting the calcofluor staining experiments that showed uniform staining (SI Appendix, Fig. S7B). Interestingly, suppressor mutant cells appeared smaller when treated with calcineurin inhibitors despite being largely restored to wild-type morphology, suggesting some fitness cost under these conditions. To validate this, we measured the area of the cells in each case from brightfield images as a proxy for cell size. As expected, wild-type cells showed a significantly larger area when treated with FK506 and CsA due to the formation of chains of cells (Fig. 4C). Surprisingly, cells with suppressor mutations harbored 10 to 20% reduced cell area even without calcineurin inhibition. This reduction became evident when cells were grown in the presence of calcineurin inhibitors and cells exhibited an approximately 50% reduction in cell area on average in each mutant analyzed. This analysis suggests either an earlier onset of cytokinesis in these cells or a compromised cell wall that does not support growth to maximum size in cells with suppressor mutations. Because the effect on cell size was more pronounced upon calcineurin inhibition, we propose that calcineurin directly regulates cytokinesis in these mutants, and this role can be bypassed through mutations in key regulators of this process.

### Chs6, Cts1 and Imp2 form an inter-dependent network

Next, we tested whether Chs6, Cts1, and Imp2 are dependent on each other for their role in cytokinesis. To study this, we deleted these genes individually in the wild-type and suppressors with mutations in each of the three genes. We hypothesize that if these proteins function in a complex, removing any component will render the suppressor mutation in other partners ineffective. As expected, the deletion of *cts1* in the wild-type strain rendered cells thermosensitive, similar to previous findings (Fig. 4D) (20). Additionally, the *imp2*Δ mutant also exhibited a thermosensitive phenotype with poor growth even at 25°C suggesting severe defects in this mutant. The *chs6*Δ strain did not have a thermosensitive phenotype on its own but exhibited hypersensitivity to the presence of FK506 and CsA at 30°C.

The deletion of these genes in respective suppressors (for example, *cts1*Δ in D3*^CTS1^*) abolished the suppressor phenotypes further supporting that mutations in these genes are responsible for the calcineurin bypass for growth (Fig. 4D). Interestingly, deletion of *cts1* or *imp2* in suppressors with a mutation in *CHS6* rendered them inviable in the presence of FK506 and CsA suggesting that Chs6 requires both Imp2 and Cts1 for its function. Similarly, the deletion of *cts1* in a suppressor with *IMP2* mutation or vice versa led to a complete cessation of growth in the presence of FK506 and CsA revealing that Imp2 and Cts1 are functionally dependent on each other. Interestingly, the deletion of *chs6* in the *CTS1* suppressor showed little hypersensitivity to FK506 and CsA at 30°C and 37°C, whereas it had no effect in the *IMP2* suppressor (Fig. 4D). Combined, these results revealed that Imp2 and Cts1 are inter-dependent on each and Chs6 is dependent on both Imp2 and Cts1. However, Imp2 does not require Chs6 and Cts1 is partially dependent on Chs6. We hypothesize that this independent phenotype of Imp2 and Cts1 could be due to the presence of Chs8, which also belongs to the same class of chitin synthases and might to some extent functionally replace Chs6 when it is absent. To this end, AlphaFold 3 modeling of Chs8 revealed not only a highly similar structure to Chs6 but also predicted the formation of a heterodimer between Chs6 and Chs8 (SI Appendix, Fig. S6F-G). Furthermore, we were unable to obtain a *chs6*Δ *chs8*Δ deletion mutant suggesting that a double deletion mutant may be inviable.

## Discussion

In this study, we defined mechanisms of calcineurin signaling enabling cytokinesis during thermal stress in *C. deneoformans*. To this end, we isolated mutations that bypass the requirement for calcineurin function during high temperature growth, and whole genome sequencing was then performed to identify the causative mutations. Our analysis revealed that proteins involved in septation play a crucial role in bypassing calcineurin functions during high temperature growth. Interestingly, these mutations did not restore sexual reproduction in the absence of calcineurin suggesting a specific role for these proteins in calcineurin-dependent cytokinesis that is essential for high temperature growth.

The genes encoding three functionally related proteins, namely Chs6, Cts1, and Imp2, were repeatedly mutated resulting in amino acid substitutions in multiple independent isolates. Orthologs of these proteins (Chs2/Chs6, Inn1/Cts1, and Hof1/Imp2) are known to be involved in cytokinesis as part of the mitotic exit network in *S. cerevisiae* (26, 37). Chs2, Inn1, and Hof1 form the Ingression Progression Complex (IPC) along with another protein Cyk3 (41, 42). Cyk3 does not have a clear ortholog in *Cryptococcus* species and it is possible that it is absent in this species and just three proteins comprise the IPC. This complex is primarily responsible for the formation of primary septa during cell division and operates in parallel with the actomyosin ring to complete cytokinesis. Hof1 is an F-BAR and SH3 domain protein that directly interacts with septins and localizes at the site of cell division during cytokinesis where it recruits Inn1 through its SH3 domain. Hof1 also directly interacts with Chs2 and provides a platform for Chs2 localization during cell division. This facilitates interaction between the C2-domain of Inn1 leading to activation of Chs2 activity resulting in chitin synthesis and primary septa formation (26, 28, 37, 38, 40, 43). Furthermore, Hof1 association with the actomyosin ring was found to be significantly impacted by phosphorylation events (47, 48). Our data revealed that calcineurin dephosphorylates the Hof1 ortholog Imp2 in *C. deneoformans* and thus might be regulating its localization to the actomyosin ring.

We hypothesize that Chs6, Cts1, and Imp2 form a complex in *C. deneoformans*, similar to their orthologs in *S. cerevisiae*, where Imp2 forms the scaffold for recruitment of Cts1 and Chs6 (Fig. 5). The association of Imp2 with either Cts1 or Chs2 or both is impacted by phosphorylation status, which in turn is regulated by calcineurin. Dephosphorylation by calcineurin at 37°C promotes the formation of this complex leading to activation of Chs6 and primary septa formation, which results in successful cytokinesis. In the absence of calcineurin activity, Chs6 is not activated, and thus primary septa fail to form resulting in cytokinesis failure. The suppressor mutations bypass the requirement for calcineurin either by promoting interactions between these proteins or by directly activating the chitin synthesis activity of Chs6. This results in primary septa formation and cytokinesis independent of calcineurin. Indeed, a previous study in *S. cerevisiae* identified dominant mutations in Chs2 that can bypass the requirement for Inn1 by constitutively activating Chs2 (26).

**Fig. 5.**
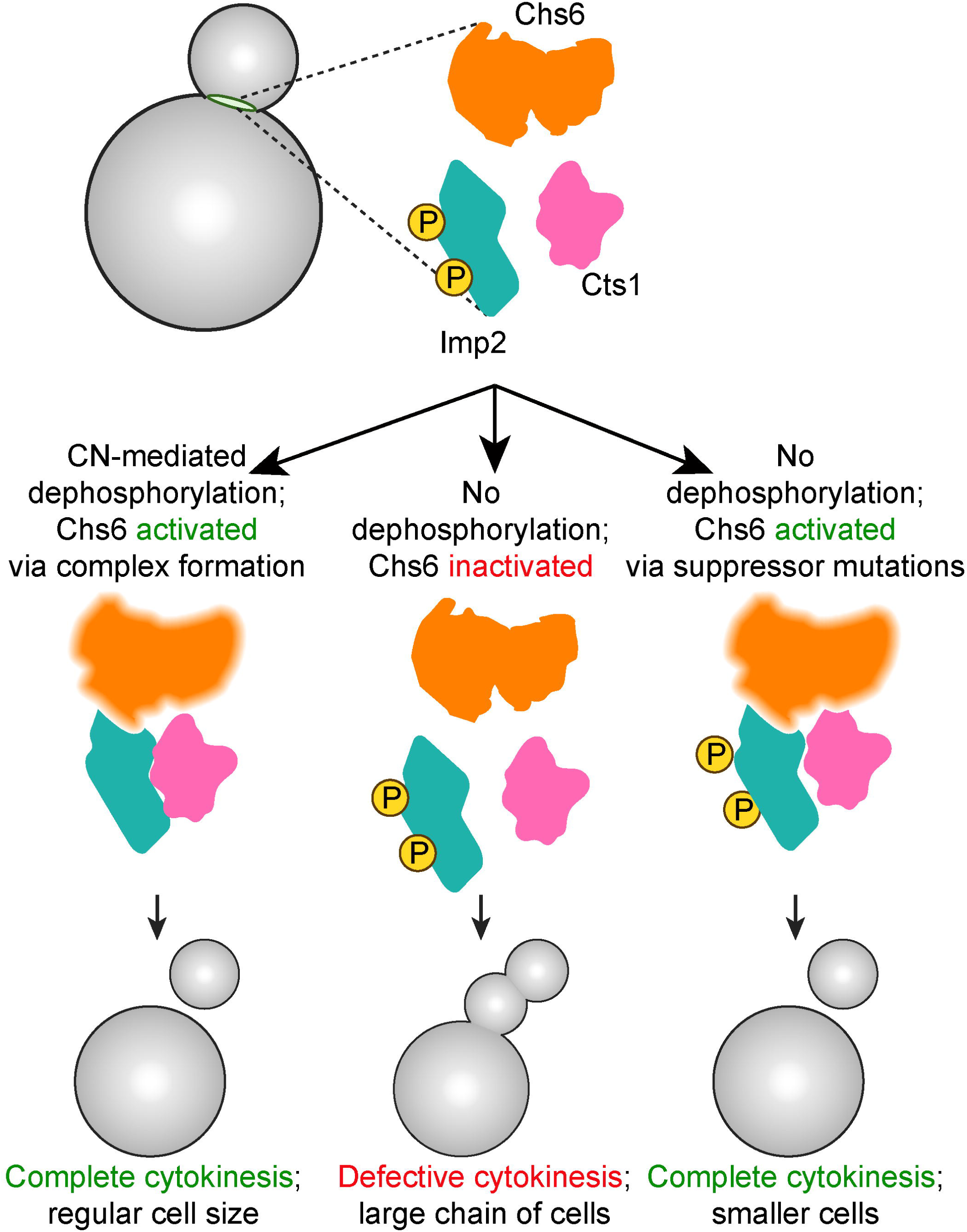
A model depicting the control of cytokinesis by calcineurin. A cartoon model depicting the impact of calcineurin-mediated dephosphorylation as well as dominant suppressor mutations on cytokinesis is presented. We hypothesize that dominant mutations in either Chs6, Cts1, or Imp2 activate Chs6 activity resulting in cytokinesis.

Homologs of Cts1 and Imp2 also play an important role in contractile ring formation in *S. pombe* suggesting a conserved role for these proteins (27, 29). Calcineurin has been shown to directly regulate the function of Cdc15/Imp2 during contractile ring formation in *S. pombe* (49–51). Our phosphoproteome analysis provides evidence that Imp2 is a substrate of calcineurin at high temperature. Thus, calcineurin might directly regulate Imp2 during cytokinesis. These findings suggest that the role of calcineurin in cytokinesis may be highly conserved across species. Furthermore, this role may be temperature-dependent because we do not observe the same cellular defects at 25°C in the absence of calcineurin. We hypothesize that Imp2 may be either dephosphorylated by a different phosphatase at 25°C or it is specifically phosphorylated by a kinase at 37°C, which then requires dephosphorylation by calcineurin. These aspects of the hypothesized network will require further investigation to resolve.

We also hypothesize that mutation in Sho1 and loss of Cdc42 observed in our screen also function in the same process. The Sho1 ortholog was found to interact with Hof1, Inn1, and Cyk3 and promote cytokinesis in *S. cerevisiae* in a similar fashion to IPC (52). Similarly, the inactivation of Cdc42 was found to promote the formation of secondary septa and restore cytokinesis defects observed in the *cyk3*Δ *hof1*Δ mutant in *S. cerevisiae* (53, 54). These findings suggest that not only the core components of IPC but also the other components of cytokinesis machinery and their functions might be conserved between *C. deneoformans* and *S. cerevisiae*. Whether these components are conserved among other budding yeasts, specifically fungal pathogens such as *C. albicans*, and function in a similar fashion requires a better understanding of cytokinesis machinery in these yeasts.

## Acknowledgments

We thank Audrey R. Odom John, currently the Stanley Plotkin Chair of Pediatric Infectious Diseases and Chief of the Division of Pediatric Infectious Diseases at the Children’s Hospital of Philadelphia, who as an undergraduate researcher in the lab between 1994-1996 froze suppressor isolates that were cross-resistant to FK506 and CsA as well as for critical reading of the manuscript. We are grateful to Professor Kenichi Yokoyama, Professor Seok-Yong Lee, and Dr. Zhenning Ren, Department of Biochemistry, Duke University School of Medicine for helpful discussions, suggestions, and critical reading of the manuscript. We also thank Prof. Lukasz Kozubowski, Clemson University, and Dr. Erica Washington, Duke University Medical Center for providing feedback on the manuscript. We also thank Dr. Zhuyun Bian in our lab and Justin Gladman, R&D Engineer at the Shared Materials Instrumentation Facility for assistance with the scanning electron microscopy image acquisition. We thank Dr. Erik Soderblom, Greg Waitt, and Tricia Ho at the Duke University Proteomics and Metabolomics Core Facility for phosphoproteome as well as Cna1 immunoprecipitation data acquisition. We also thank Dr. Devi Swain Lenz and the Duke University Sequencing and Genomic Technologies (SGT) Core Facility for Illumina sequencing. We kindly acknowledge Dr. Connie Nichols and Professor Andrew Alspaugh, Duke University School of Medicine for providing the *C. neoformans cdc42*Δ strains and helpful discussions. This work was supported by NIH/NIAID R01 awards AI039115-27, AI050113-20, and AI172451-02 awarded to JH. JH is also a Co-Director and Fellow of the CIFAR program Fungal Kingdom: Threats & Opportunities.

## Materials and methods

### Strains and media

*Cryptococcus deneoformans* (previously known as *C. neoformans* var. *neoformans*) reference strains JEC20**a** and JEC21α were used for experiments conducted in this study (55, 56). The strains were cultured in YPD (1% yeast extract, 2% peptone, and 2% glucose) liquid or YPD agar media at 25°C or 37°C depending on the experimental requirements. Yeast Nitrogen Base without amino acids supplemented with 2% glucose was used to make standard defined (SD) media without uracil and/or adenine (SD-ura-ade or SD-ura or SD-ade). For calcineurin inhibition conditions, calcineurin inhibitors FK506 and CsA were added to the YPD agar media at a final concentration of 1 µg/ml and 50 µg/ml, respectively. Gene deletion mutants and ectopic tagged strains were generated via homologous recombination using CRISPR following the TRACE protocol (57) and transformants were selected on YPD agar media supplemented with either 100 µg/ml nourseothricin or 200 µg/ml G-418. The strains generated and used in this study are presented in SI Appendix, Table S3 and primers used in this study are presented in SI Appendix, Table S4.

### Isolation of calcineurin suppressors

Spontaneous genetic mutations that suppress calcineurin’s role for high temperature growth were isolated in either a wild-type strain background (JEC20 and JEC21) or their respective *cnb1*Δ deletion mutants (JEC20 *cnb1*Δ and JEC21 *cnb1*Δ) (Fig. 1A). In each case, the strains were first streaked to obtain single colonies on YPD media at 25°C. Next, 12 to 15 independent single colonies were inoculated for each strain in 5 ml YPD media and grown overnight at 25°C. An equal number of cells, equivalent to 1 OD, were plated on either YPD agar containing both FK506 and CsA for wild-type strains or YPD agar media for *cnb1*Δ strains and incubated at 37°C. The plates were incubated for 4-5 days to allow the growth of mutant strains that exhibited robust growth in the restrictive conditions. For each plate, 4 colonies were again streaked on respective restrictive conditions and verified for robust growth. Out of these, only one colony was then used for further analysis as an independent genetic mutant for calcineurin suppression, and a total of 52 suppressors were isolated through this approach. Additionally, 12 suppressors were isolated in a previous study as FK506-resistant colonies that were cross-resistant to CsA (5).

### Whole-genome sequencing and variant calling

A total of 64 calcineurin suppressors along with JEC20 *cnb1*Δ and JEC21 *cnb1*Δ mutant strains were subjected to Illumina whole genome sequencing. Each strain was grown overnight in 5 ml YPD liquid media. Cells were harvested by centrifugation at room temperature and the cell pellet was frozen at −80C. The cells were lyophilized, and DNA extraction was performed using the MasterPure yeast DNA purification kit (Biosearch Technologies). The isolated DNA was checked for quality via agarose gel electrophoresis and NanoDrop and was quantified using Qubit. The DNA isolated was then submitted to Duke Sequencing and Genomic Technologies core for sequencing on the Illumina Novaseq platform.

The sequencing reads obtained were checked for quality and mapped to *C. deneoformans* reference JEC21 genome available on FungiDB (https://fungidb.org/fungidb/app) using Bowtie2 with default parameters and allowing every read to map only once. The BAM files containing mapped reads were converted to TDF using IGVtools and visualized in IGV for the genome coverage plots and to identify events of aneuploidy. In parallel, the BAM files were analyzed using Geneious Prime for variant identification in each case. Single Nucleotide Polymorphism (SNP) events that were covered with at least 60 reads and were present in 90% of covering reads were scored as variants and were considered for analysis. Among these, SNPs that were shared between the suppressor isolates and *cnb1*Δ mutants were not considered for further analysis. This analysis resulted in the identification of only one SNP in the protein-coding regions per isolate for the majority of the isolates. For isolates that did not present any SNP in their protein-coding regions and did not harbor any aneuploidy events, SNPs present in the introns were analyzed and considered. Combined together, we were able to identify a SNP or aneuploidy event in each isolate with the majority of them harboring only one genetic change. The mutations identified are summarized in Table 1 and Supplementary Dataset S1.

### Mating and cell-cell fusion assays

Mating assays to determine the impact of suppressor mutations on sexual reproduction were performed on MS media at room temperature. The parental strains of opposite mating types for each cross were grown on YPD agar plates for 2 days. Approximately equal number of cells for each parent were harvested from the plate and mixed together in 200 µl of purified and autoclaved water. The cell suspension was mixed thoroughly by vortexing and 3 µl of the cell suspension was spotted on the MS media plates and incubated for 2 weeks at room temperature. To assess the impact of calcineurin inhibition, FK506 was added to the MS media at a final concentration of 1 µg/ml, and assays were done in parallel for each cross.

Cell-cell fusion assays to generate diploids were done by crossing the selected suppressors with the opposite mating type partner and were selected by auxotrophic selection markers (Figure 3B). To obtain uracil auxotrophs in the selected calcineurin suppressors and a wild-type strain, 1 OD equivalent cells were plated on SD media supplemented with 5-FOA and incubated at 30°C for 7 days. The resistant colonies were streaked on the 5-FOA-containing media as well as SD media lacking uracil (SD-ura) to confirm the auxotrophs. Genomic DNA was prepared from the auxotrophs, *URA5* gene was PCR amplified and sequenced via Plasmidsaurus to confirm the mutation in the *URA5* gene. One confirmed auxotroph for each suppressor was then used for cell-cell fusion assays and crossed with an adenine auxotrophic strain of the opposite mating type. Cells were mixed and spotted on the MS media similar to as described for mating assays and incubated in the MS media for 48 hours to allow cell-cell fusion. Cells were then scraped from the MS media, resuspended in water, diluted serially, spread on SD-ura-ade plates, and incubated at 37°C for 4 to 5 days for selecting prototrophic diploids. The isolated colonies were further streaked on SD-ura-ade media and then analyzed by flow cytometry to confirm the ploidy, by PCR analysis to confirm the presence of both mating type loci in their genome, growth SD-ura-ade media and their ability to self-filament in the MS media at room temperature. For each suppressor and the wild-type strain, 8 confirmed diploids were analyzed for their ability to grow in the presence of calcineurin inhibitors, FK506 and CsA, at 37°C and scored for dominant versus recessive nature of the suppressor mutation.

### Cell morphology and size analysis

To study the impact of calcineurin inhibition on wild-type and suppressors, the cells were grown overnight in the YPD media. The cultures were diluted in the fresh media starting with 0.2 OD and were grown in the presence or absence of 1 µg/ml FK506 and 50 µg/ml CsA for 20 hours. Cellular morphology was observed using a Zeiss AxioVision microscope at 100 X magnification in each case. Cells were stained with calcofluor white to stain the chitin as described previously (58) and to assess the site of septation.

Cell size was determined as the surface area for each cell in the images. The acquired brightfield images were processed using YeastSpotter (59) to obtain the segmentation images compatible with ImageJ/Fiji. The processed images were used as the input in Fiji (60) to quantify the surface area for each cell. For wild-type cells that form chains of cells, the surface area was manually calculated for each cell by combining individual values obtained from Fiji. The measured values were plotted using GraphPad Prism.

### Chitin-Chitosan measurement assays

Chitin and chitosan levels were quantified using 3-methyl-2-benzothiazolone hydrazine hydrochloride (MBTH)-based assay, as previously described (58). Briefly, 20 ml cultures were grown in triplicates in the same manner as described for cell morphology determination assays where one set was treated with FK506 and CsA whereas one set was untreated. Cells were harvested, lyophilized, and weighed to determine dry biomass. Pelleted cells were resuspended in 10 mL of 6% KOH and incubated at 80°C for 30 minutes with intermittent vortexing to remove background molecules that may potentially react with MBTH. After four PBS washes to remove residual alkali, the cells were resuspended in PBS at 10 mg/ml. Each sample was then split into two parts. One part (Part A) was directly subjected to chitosan quantification using the MBTH assay (58). The second part (Part B) was treated with an equal volume of 1M HCl and incubated at 110°C for 2 hours to fully deacetylate cell wall chitin into chitosan, which was then measured using the MBTH assay. Chitosan content was expressed as nanomoles of glucosamine per unit dry weight of cell wall material (Figure S7B). Chitin levels were determined by subtracting the chitosan measured in Part A from the total chitosan measured in Part B. The glucosamine content in Part A reflected the native chitosan level in the sample.

### Phosphoproteome sample preparation and analysis

The wild-type strain JEC21 and a calcineurin mutant JEC21 *cnb1*Δ were grown in 50 ml liquid YPD for 24 hours at 25°C in triplicates. The culture was then split into two equal halves of 25 ml cultures with 25 OD equivalent cells in each. One set was further incubated at 25°C for 2 hours whereas the second set was subjected to heat stress at 37°C for 2 hours. Cells were harvested from all samples by centrifugation and immediately frozen at −80°C. All 12 samples were then used for phosphoproteome analysis at the Duke University Proteomics and Metabolomics Core Facility.

Samples were supplemented with 100 µl of 8 M urea in 50 mM ammonium bicarbonate before protein extraction using a bead beater (3 rounds at 10 sec). Following quantification by Bradford assay, 200 µg of protein from each sample was used for further analysis. Each sample was spiked with 5 or 10 pmol bovine casein as an internal quality control standard. Next, the samples were reduced for 30 min at 32°C, alkylated with 20 mM iodoacetamide for 30 min at room temperature, then supplemented with a final concentration of 1.2% phosphoric acid and 1342 μl of S-Trap (Protifi) binding buffer (90% MeOH/100mM TEAB). Proteins were trapped on the S-Trap micro cartridge, digested using 40 ng/μl sequencing grade trypsin (Promega) for 1 hr at 47°C, and eluted using 50 mM TEAB, followed by 0.2% FA, and lastly using 50% ACN/0.2% FA. All samples were then lyophilized to dryness. The phosphopeptide samples were resuspended in 80% acetonitrile and 1% TFA prior to TiO2 enrichment. Each sample was subjected to complex TiOx enrichment using GL biosciences TiO2 tips and manufacturer-recommended protocols. Eluted phosphopeptides were then subjected to C18 stage tip cleanup. All samples were frozen and lyophilized.

Quantitative LC/MS/MS was performed on 4 µl of each sample using a Vanquish Neo UPLC system coupled to a Thermo Orbitrap Astral high-resolution accurate mass tandem mass spectrometer. Briefly, the sample was first trapped on a Symmetry C18 20 mm × 180 μm trapping column (5 μl/min at 99.9/0.1 v/v water/acetonitrile), after which the analytical separation was performed using a 1.5 μm EvoSep 150 µm ID x 8cm performance (EveoSep) column with a 30 min gradient of 5 to 30% acetonitrile with 0.1% formic acid at a flow rate of 500 nanoliters/minute (nl/min) with a column temperature of 50°C. Data collection on the Orbitrap Astral mass spectrometer was performed in a data-independent acquisition (DIA) mode of acquisition with r=240,000 (@ m/z 200) full MS scan from m/z 380-1080 with a target AGC value of 4e5 ions. Fixed DIA windows of 5 m/z from m/z 380 to 1080 DIA MS/MS scans were acquired in the Astral with a target AGC value of 5e4 and a max fill time of 8 ms. HCD collision energy setting of 27% was used for all MS2 scans. The total analysis cycle time for each sample injection was approximately 45 min.

Following 16 total UPLC-MS/MS analyses (12 experimental samples and 4 internal quality control samples), data were imported into Spectronaut (Biognosis), and individual LC-MS data files were aligned based on the accurate mass and retention time of detected precursor and fragment ions. Relative peptide abundance was measured based on MS2 fragment ions of selected ion chromatograms of the aligned features across all runs. The MS/MS data was searched against a *C. deneoformans* JEC21 database, a common contaminant/spiked protein database (bovine albumin, bovine casein, yeast ADH, etc.), and an equal number of reversed sequence “decoys” for false discovery rate determination. A library was generated with only the LC-MS data files collected within this study. Database search parameters included fixed modification on Cys (carbamidomethyl) and variable modification on Met (oxidation), Protein N-term (acetyl), and Ser/Thr/Tyr (phos). Full trypsin enzyme rules were used along with 10ppm mass tolerances on precursor ions and 20ppm on product ions. Spectral annotation was set at a maximum 1% peptide false discovery rate based on q-value calculations. Note that peptide homology was addressed using razor rules in which a peptide matched to multiple different proteins was exclusively assigned to the protein that has more identified peptides.

The output raw peptide intensity values from the Spectronaut detection software were noted. At this stage, any peptide that was not detected a minimum of 2 times across all of the samples and not detected in at least 50% of one of the biological groups was removed from further analysis. For remaining peptides, any missing data values were imputed using the following rules: 1) if less than 50% of signals were missing within a group, a randomized value near the average of the remaining values was calculated, or 2) if >50% of the signals were missing within the group, a randomized intensity within the bottom 1% of the detectable signals was used. Next, non-phosphorylated peptides and phosphopeptides which did not pass a 75% confidence of localization were removed from the analysis. The remaining phosphopeptides were then subjected to a robust mean normalization in which the highest and lowest 10% of the phosphopeptide signals were ignored and the average value of the remaining phosphopeptides was used as the normalized factor across all samples. We then summed all of the same precursor states for the same phosphopeptide (i.e. different charge states) into a single modified phosphopeptide value and calculated the % coefficient of variation across each group. Following database searching and peptide scoring using a library-based Spectronaut search, the data was annotated at a 1% peptide false discovery rate. A total of 14,484 phosphopeptides were identified in the dataset corresponding to 2,040 phosphoproteins.

The normalized values were then used for analysis to calculate the fold-change between various groups and significantly enriched peptides in each group were identified using DEP2 (61). Analysis was done using R scripts to identify the phosphopeptides that were significantly enriched in the JEC21 *cnb1*Δ mutant at 37°C specifically. All peptides that exhibited at least a 1.5-log2-fold difference and a p-value < 0.05 were considered for the analysis. All sites in the calcineurin suppressor proteins were identified from this dataset and marked in the volcano plots presented in Figure 3D-E.

### Fluorescence microscopy

Cells expressing Cna1-GFP and Cdc10-mCherry (VYD371) were grown overnight in 5 ml of YPD liquid. The next day, 2 OD cells were transferred to 5 ml of fresh YPD media and were grown for 3 hours. Following that growth, cells were harvested and washed twice with 1 ml of purified water each. Finally, cells were resuspended in 200 µl of YNB media with 2% glucose. From this, 5 µl of cell suspension was spotted onto a 2% agarose slab prepared in YNB+2% glucose media, which was then inverted into a glass bottom MatTek dish. The cells were then imaged using the DeltaVision microscope system at the Duke Light Microscope Core Facility. The images were captured using mCherry and GFP filters for Cdc10-mCherry and Cna1-GFP, respectively with a time interval of 2-4 minutes each with the microscope chamber temperature set at 37°C. The images were then processed using SoftWoRx software, v 6.1, connected to the microscope and ImageJ. The processed images were assembled using Adobe Photoshop for presentation purposes.

### Cna1-GFP immunoprecipitation

Strain expressing Cna1-GFP were grown in 5 ml YPD media overnight in triplicates. Next day, each triplicate sample was transferred to four new 50 ml fresh cultures with starting OD600 of 0.01 each (4 sets of triplicates i.e. 12 cultures in total) and grown for 20 hours at 25°C with vigorous shaking. One set was further incubated at 25°C, second set was moved to 37°C, third set was treated with FK506 at 25°C and the fourth set was treated with FK506 and moved to 37°C. Each of the sets was further grown for 5 hours before cell harvesting. Harvested cells were lyophilized and lysed by bead-beating in the RIPA buffer. The lysates were purified, and protein was quantified. One mg of protein per sample was used as input for immunoprecipitation, which was performed using Chromotek GFP-TRAP beads using the manufacturer’s guidelines. The bound proteins were eluted in 100 µl of elution buffer (1% SDS; 10mM DTT) at 90°C for 10 mins. A fraction of the eluted sample (10 µl) was used for validation by Western blotting and probed for GFP whereas the rest was submitted for mass-spectrometry analysis at the Duke University Proteomics and Metabolomics Core Facility.

### AlphaFold modeling

AlphaFold modeling was done using AlphaFold3 available online (https://alphafoldserver.com/). The models were generated in various combinations and configurations for Chs6, Cts1, Imp2 and Chs8. For the hexameric configuration model presented in Figure 4A (Chs6 x2; Imp2 x2; Cts1 x2), a truncated version of Cts1 lacking amino acids 309-708 was used to account for 5000 amino acid limits for AlphaFold modeling. This region of Cts1 does not harbor any suppressor mutation or a conserved domain and thus was selected for truncation in the model. For each prediction, 5 different models were obtained from the AlphaFold server and only the model_0.cif were used for analysis and presentation in this study. All generated models were processed using ChimeraX 1.9 for presentation purposes (62, 63).

### Data availability

The whole genome sequencing data for all the suppressors and calcineurin mutant strains is available via NCBI BioProject PRJNA1202532. The mass spectrometry proteomics data have been deposited to the ProteomeXchange Consortium (64) via the PRIDE (65) partner repository with the dataset identifiers PXD060167 (Cna1-GFP IP data) and PXD059891 (phosphoproteome data).

## References

1. R. Rajasingham et al., The global burden of HIV-associated cryptococcal infection in adults in 2020: A modelling analysis. Lancet Infect Dis 22, 1748–1755 (2022).

2. K. J. Kwon-Chung et al., *Cryptococcus neoformans* and *Cryptococcus gattii*, the Etiologic Agents of Cryptococcosis. Cold Spring Harb Perspect Med 4, a019760–a019760 (2014).

3. Y. Zhao, J. Lin, Y. Fan, X. Lin, Life cycle of *Cryptococcus neoformans*. Annu Rev Microbiol 73, 17–42 (2019).

4. D. S. Fox et al., Calcineurin regulatory subunit is essential for virulence and mediates interactions with FKBP12-FK506 in *Cryptococcus neoformans*. Mol Microbiol 39, 835–849 (2001).

5. A. Odom et al., Calcineurin is required for virulence of *Cryptococcus neoformans*. EMBO J 16, 2576–2589 (1997).

6. I. Ulengin-Talkish, M. S. Cyert, A cellular atlas of calcineurin signaling. Biochim Biophys Acta Mol Cell Res 1870, 119366 (2023).

7. V. Yadav, J. Heitman, Calcineurin: The Achilles’ heel of fungal pathogens. PLoS Pathogens 19, e1011445 (2023).

8. H. S. Park, S. C. Lee, M. E. Cardenas, J. Heitman, Calcium-calmodulin-calcineurin signaling: A globally conserved virulence cascade in eukaryotic microbial pathogens. Cell Host Microbe 26, 453–462 (2019).

9. W. J. Steinbach, J. L. Reedy, R. A. Cramer, Jr., J. R. Perfect, J. Heitman, Harnessing calcineurin as a novel anti-infective agent against invasive fungal infections. Nat Rev Microbiol 5, 418–430 (2007).

10. E. W. Chow et al., Elucidation of the calcineurin-Crz1 stress response transcriptional network in the human fungal pathogen *Cryptococcus neoformans*. PLoS Genetics 13, e1006667 (2017).

11. C. S. Hemenway, J. Heitman, Calcineurin: Structure, function, and inhibition. Cell Biochem Biophys 30, 115–151 (1999).

12. D. P. Matheos, T. J. Kingsbury, U. S. Ahsan, K. W. Cunningham, Tcn1p/Crz1p, a calcineurin-dependent transcription factor that differentially regulates gene expression in *Saccharomyces cerevisiae*. Genes Dev 11, 3445–3458 (1997).

13. J. Roy, M. S. Cyert, Identifying new substrates and functions for an old enzyme: Calcineurin. Cold Spring Harb Perspect Biol 12, a035436 (2020).

14. S. Ho et al., The mechanism of action of cyclosporin A and FK506. Clin Immunol Immunopathol 80, S40–45 (1996).

15. L. Kozubowski, E. F. Aboobakar, M. E. Cardenas, J. Heitman, Calcineurin colocalizes with P-bodies and stress granules during thermal stress in *Cryptococcus neoformans*. Eukaryot Cell 10, 1396–1402 (2011).

16. L. Kozubowski, J. W. Thompson, M. E. Cardenas, M. A. Moseley, J. Heitman, Association of calcineurin with the COPI protein Sec28 and the COPII protein Sec13 revealed by quantitative proteomics. PLoS One 6, e25280 (2011).

17. H. S. Park et al., Calcineurin targets involved in stress survival and fungal virulence. PLoS Pathogens 12, e1005873 (2016).

18. M. E. Cardenas, R. S. Muir, T. Breuder, J. Heitman, Targets of immunophilin-immunosuppressant complexes are distinct highly conserved regions of calcineurin A. EMBO J 14, 2772–2783 (1995).

19. M. C. Cruz, D. S. Fox, J. Heitman, Calcineurin is required for hyphal elongation during mating and haploid fruiting in *Cryptococcus neoformans*. EMBO J 20, 1020–1032 (2001).

20. D. S. Fox, G. M. Cox, J. Heitman, Phospholipid-binding protein Cts1 controls septation and functions coordinately with calcineurin in *Cryptococcus neoformans*. Eukaryot Cell 2, 1025–1035 (2003).

21. E. F. Aboobakar, X. Wang, J. Heitman, L. Kozubowski, The C2 domain protein Cts1 functions in the calcineurin signaling circuit during high-temperature stress responses in *Cryptococcus neoformans*. Eukaryot Cell 10, 1714–1723 (2011).

22. E. R. Ballou, C. B. Nichols, K. J. Miglia, L. Kozubowski, J. A. Alspaugh, Two CDC42 paralogues modulate *Cryptococcus neoformans* thermotolerance and morphogenesis under host physiological conditions. Mol Microbiol 75, 763–780 (2010).

23. Y. C. Chang, A. Khanal Lamichhane, K. J. Kwon-Chung, *Cryptococcus neoformans*, unlike *Candida albicans*, forms aneuploid clones directly from uninucleated cells under fluconazole stress. mBio 9, 01290–01218 (2018).

24. F. Yang et al., Adaptation to fluconazole via aneuploidy enables cross-adaptation to amphotericin B and flucytosine in *Cryptococcus neoformans*. Microbiol Spectr 9, e0072321 (2021).

25. T. Kamei et al., Interaction of Bnr1p with a novel Src homology 3 domain-containing Hof1p. Implication in cytokinesis in *Saccharomyces cerevisiae*. J Biol Chem 273, 28341–28345 (1998).

26. A. Devrekanli, M. Foltman, C. Roncero, A. Sanchez-Diaz, K. Labib, Inn1 and Cyk3 regulate chitin synthase during cytokinesis in budding yeasts. J Cell Sci 125, 5453–5466 (2012).

27. R. H. Roberts-Galbraith et al., Dephosphorylation of F-BAR Protein Cdc15 modulates its conformation and stimulates its scaffolding activity at the cell division site. Mol Cell 39, 86–99 (2010).

28. A. Sanchez-Diaz et al., Inn1 couples contraction of the actomyosin ring to membrane ingression during cytokinesis in budding yeast. Nat Cell Biol 10, 395–406 (2008).

29. A. H. Willet et al., The F-BAR Cdc15 promotes contractile ring formation through the direct recruitment of the formin Cdc12. J Cell Biol 208, 391–399 (2015).

30. M. D. Lenardon, C. A. Munro, N. A. Gow, Chitin synthesis and fungal pathogenesis. Curr Opin Microbiol 13, 416–423 (2010).

31. Y. Oh et al., Mitotic exit kinase Dbf2 directly phosphorylates chitin synthase Chs2 to regulate cytokinesis in budding yeast. Mol Biol Cell 23, 2445–2456 (2012).

32. K. Preechasuth et al., Cell wall protection by the *Candida albicans* class I chitin synthases. Fungal Genet Biol 82, 264–276 (2015).

33. C. P. Wigington et al., Systematic discovery of short linear motifs decodes calcineurin phosphatase signaling. Mol Cell 79, 342–358 e312 (2020).

34. B. L. Brauer et al., Leveraging new definitions of the LxVP SLiM to discover novel calcineurin regulators and substrates. ACS Chem Biol 14, 2672–2682 (2019).

35. L. Kozubowski, J. Heitman, Septins enforce morphogenetic events during sexual reproduction and contribute to virulence of *Cryptococcus neoformans*. Mol Microbiol 75, 658–675 (2010).

36. S. D. Singh et al., Hsp90 governs echinocandin resistance in the pathogenic yeast *Candida albicans* via calcineurin. PLoS Pathog 5, e1000532 (2009).

37. F. Meitinger et al., Targeted localization of Inn1, Cyk3 and Chs2 by the mitotic-exit network regulates cytokinesis in budding yeast. J Cell Sci 123, 1851–1861 (2010).

38. R. Nishihama et al., Role of Inn1 and its interactions with Hof1 and Cyk3 in promoting cleavage furrow and septum formation in *S. cerevisiae*. J Cell Biol 185, 995–1012 (2009).

39. S. Palani, F. Meitinger, M. E. Boehm, W. D. Lehmann, G. Pereira, Cdc14-dependent dephosphorylation of Inn1 contributes to Inn1–Cyk3 complex formation. J Cell Sci 125, 3091–3096 (2012).

40. M. Wang, R. Nishihama, M. Onishi, J. R. Pringle, Role of the Hof1–Cyk3 interaction in cleavage-furrow ingression and primary-septum formation during yeast cytokinesis. Mol Biol Cell 29, 597–609 (2018).

41. M. Foltman et al., Ingression progression complexes control extracellular matrix remodelling during cytokinesis in budding yeast. PLoS Genet 12, e1005864 (2016).

42. M. Foltman, A. Sanchez-Diaz, Central role of the actomyosin ring in coordinating cytokinesis steps in budding yeast. J Fungi 10 (2024).

43. Y. Oh, J. Schreiter, R. Nishihama, C. Wloka, E. Bi, Targeting and functional mechanisms of the cytokinesis-related F-BAR protein Hof1 during the cell cycle. Mol Biol Cell 24, 1305–1320 (2013).

44. J. Abramson et al., Accurate structure prediction of biomolecular interactions with AlphaFold 3. Nature 630, 493–500 (2024).

45. Z. Ren et al., Structural basis for inhibition and regulation of a chitin synthase from *Candida albicans*. Nat Struct Mol Biol 29, 653–664 (2022).

46. D. D. Chen et al., Structure, catalysis, chitin transport, and selective inhibition of chitin synthase. Nat Commun 14, 4776 (2023).

47. F. Meitinger et al., Phosphorylation-dependent regulation of the F-BAR protein Hof1 during cytokinesis. Genes Dev 25, 875–888 (2011).

48. F. Meitinger, S. Palani, B. Hub, G. Pereira, Dual function of the NDR-kinase Dbf2 in the regulation of the F-BAR protein Hof1 during cytokinesis. Mol Biol Cell 24, 1290–1304 (2013).

49. R. Martin-Garcia et al., Paxillin-mediated recruitment of calcineurin to the contractile ring Is required for the correct progression of cytokinesis in fission yeast. Cell Rep 25, 772–783 e774 (2018).

50. C. E. Snider, R. Bhattacharjee, M. G. Igarashi, K. L. Gould, Fission yeast paxillin contains two Cdc15 binding motifs for robust recruitment to the cytokinetic ring. Mol Biol Cell 33, br4 (2022).

51. K. Bellingham-Johnstun, B. Commer, B. Levesque, Z. L. Tyree, C. Laplante, Imp2p forms actin-dependent clusters and imparts stiffness to the contractile ring. Mol Biol Cell 33, ar145 (2022).

52. K. Labedzka et al., Sho1p connects the plasma membrane with proteins of the cytokinesis network through multiple isomeric interaction states. J Cell Sci 125, 4103–4113 (2012).

53. B. D. Atkins et al., Inhibition of Cdc42 during mitotic exit is required for cytokinesis. J Cell Biol 202, 231–240 (2013).

54. M. Onishi, N. Ko, R. Nishihama, J. R. Pringle, Distinct roles of Rho1, Cdc42, and Cyk3 in septum formation and abscission during yeast cytokinesis. J Cell Biol 202, 311–329 (2013).

55. K. J. Kwon-Chung, J. C. Edman, B. L. Wickes, Genetic association of mating types and virulence in *Cryptococcus neoformans*. Infect Immun 60, 602–605 (1992).

56. J. Heitman, B. Allen, J. A. Alspaugh, K. J. Kwon-Chung, On the origins of congenic *MAT*α and *MAT***a** strains of the pathogenic yeast *Cryptococcus neoformans*. Fungal Genet Biol 28, 1–5 (1999).

57. Y. Fan, X. Lin, Multiple applications of a transient CRISPR-Cas9 coupled with electroporation (TRACE) system in the *Cryptococcus neoformans* species complex. Genetics 208, 1357–1372 (2018).

58. B. T. Maybruck, R. Upadhya, W. C. Lam, C. A. Specht, J. K. Lodge, Fluorescence and biochemical assessment of the chitin and chitosan content of *Cryptococcus*. Methods Mol Biol 2775, 329–347 (2024).

59. A. X. Lu, T. Zarin, I. S. Hsu, A. M. Moses, YeastSpotter: accurate and parameter-free web segmentation for microscopy images of yeast cells. Bioinformatics 35, 4525–4527 (2019).

60. J. Schindelin et al., Fiji: an open-source platform for biological-image analysis. Nat Methods 9, 676–682 (2012).

61. Z. Feng, P. Fang, H. Zheng, X. Zhang, DEP2: an upgraded comprehensive analysis toolkit for quantitative proteomics data. Bioinformatics 39 (2023).

62. E. C. Meng et al., UCSF ChimeraX: Tools for structure building and analysis. Protein Sci 32, e4792 (2023).

63. E. F. Pettersen et al., UCSF ChimeraX: Structure visualization for researchers, educators, and developers. Protein Sci 30, 70–82 (2021).

64. E. W. Deutsch et al., The ProteomeXchange consortium at 10 years: 2023 update. Nucleic Acids Res 51, D1539–D1548 (2023).

65. Y. Perez-Riverol et al., The PRIDE database at 20 years: 2025 update. Nucleic Acids Res 53, D543–D553 (2025).

