## Supplementary Appendix for "Calcineurin controls the cytokinesis machinery during thermal stress in *Cryptococcus deneoformans*"

### Supplementary figures and figure legends

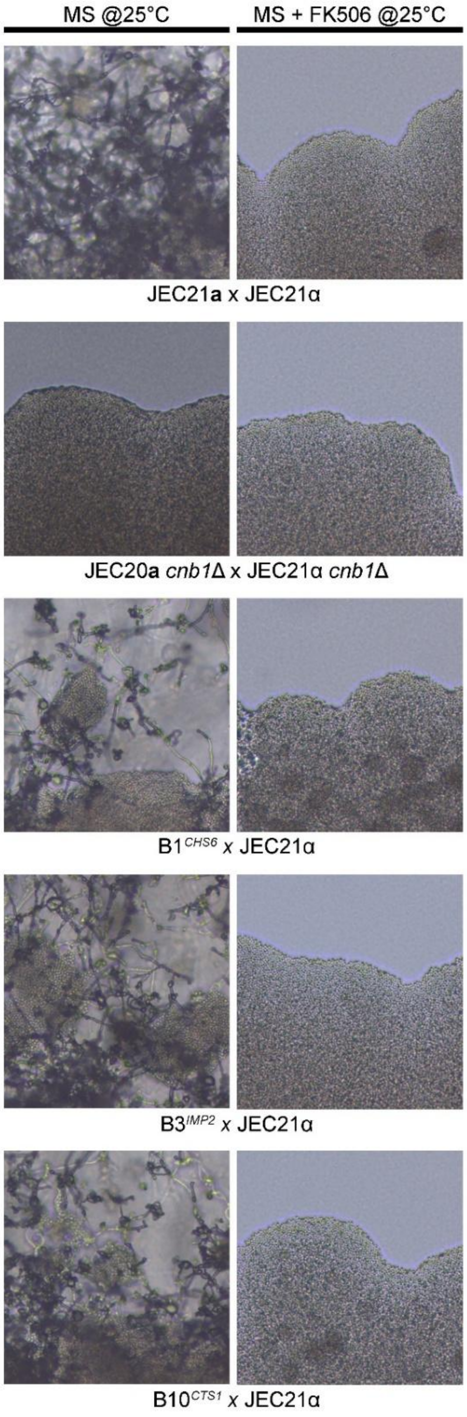


**Fig. S1. Identified mutations do not suppress calcineurin role in sexual reproduction.** Mating hyphae of crosses was imaged after 3 weeks of incubation on mating media and photographed. The wild type as well as the suppressor mutants formed hyphae and spores as expected when incubated on MS media but failed to produce hyphae in the presence of the calcineurin inhibitor FK506. The JEC21 *cnb1*Δ x JEC20 *cnb1*Δ cross is shown as a negative control in which the bilateral mutant cross failed to produce hyphae. Images were captured at 20X magnification.

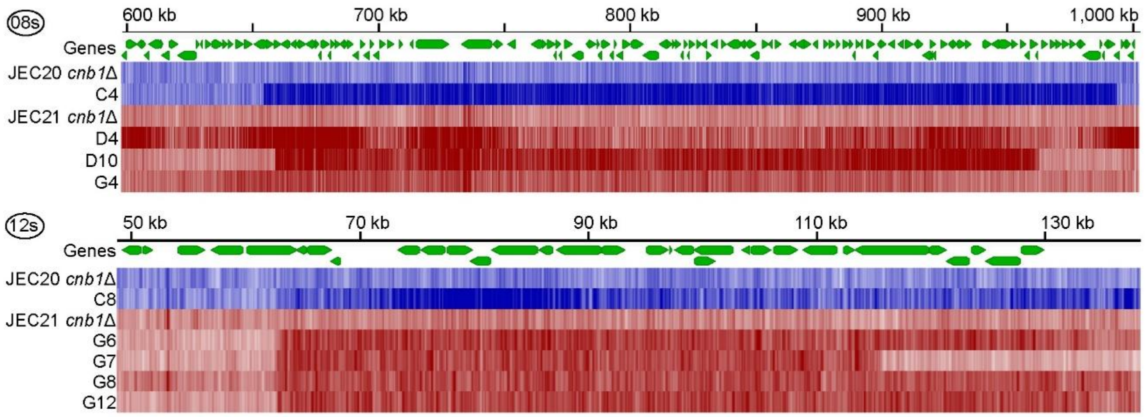


**Fig. S2. Segmental aneuploidy drives calcineurin bypass.** Genome coverage maps of two regions highlighted in Figure 2 show the aneuploidy events in chromosome 08 (08s) and chromosome 12 (12s). The list of genes present in these regions is included in the supplementary Dataset S2.

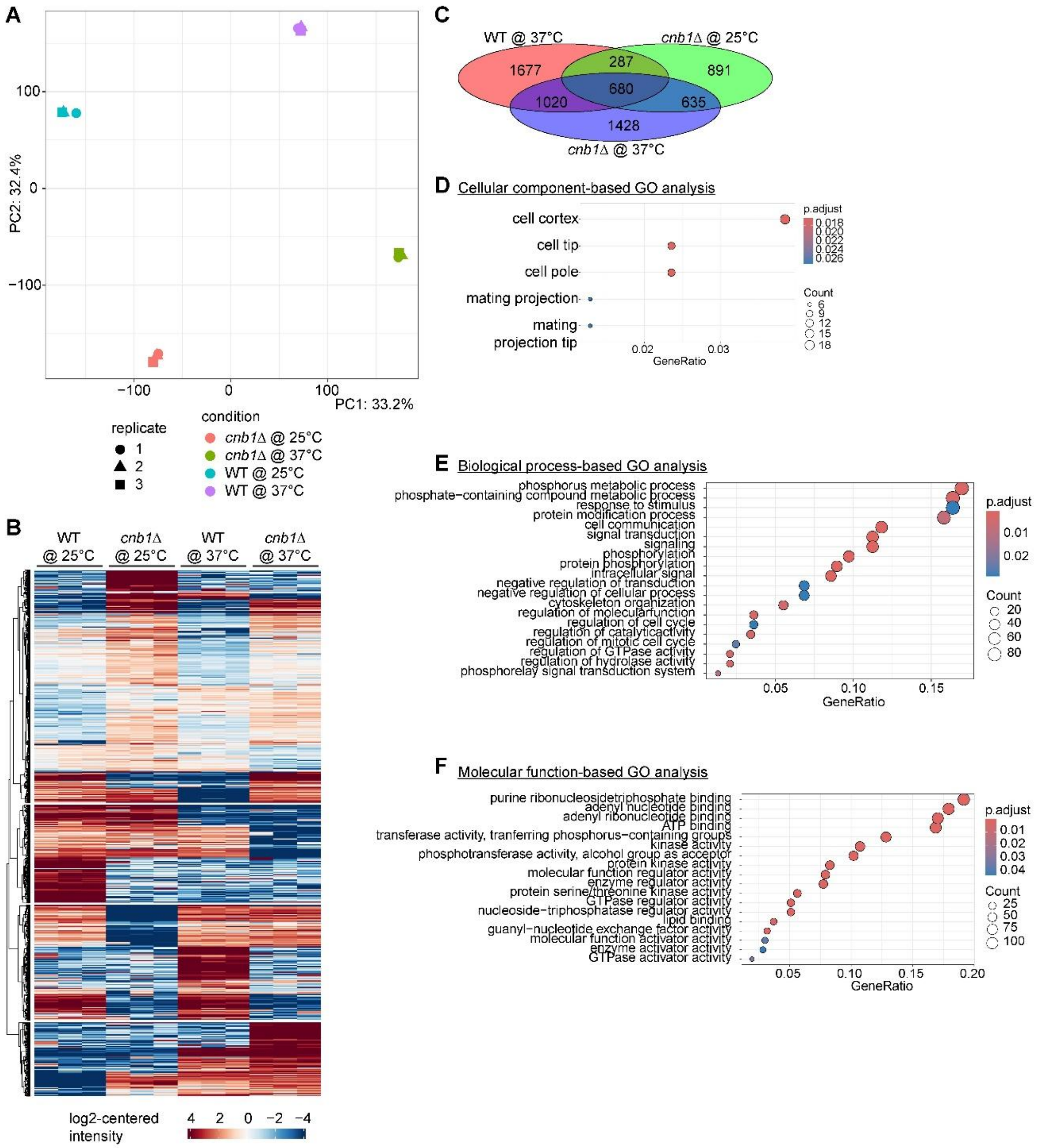


**Fig. S3. Phosphoproteome analysis for calcineurin mutants**. (A) PCA plot analysis showed clustering of three replicates for each experimental condition. (B) Heatmap analysis of all peptides showed a strong correlation of differentially phosphorylated peptides among replicates and identified condition-specific peptides. (C) A Venn diagram presenting the overlap among phosphosites that were enriched in three groups as compared to the wild type at 25°C. (D-F) Gene ontology of proteins corresponding to phosphopeptides (1428 phosphopeptides from C) that were exclusively enriched in *cnb1*Δ mutant at 37°C.

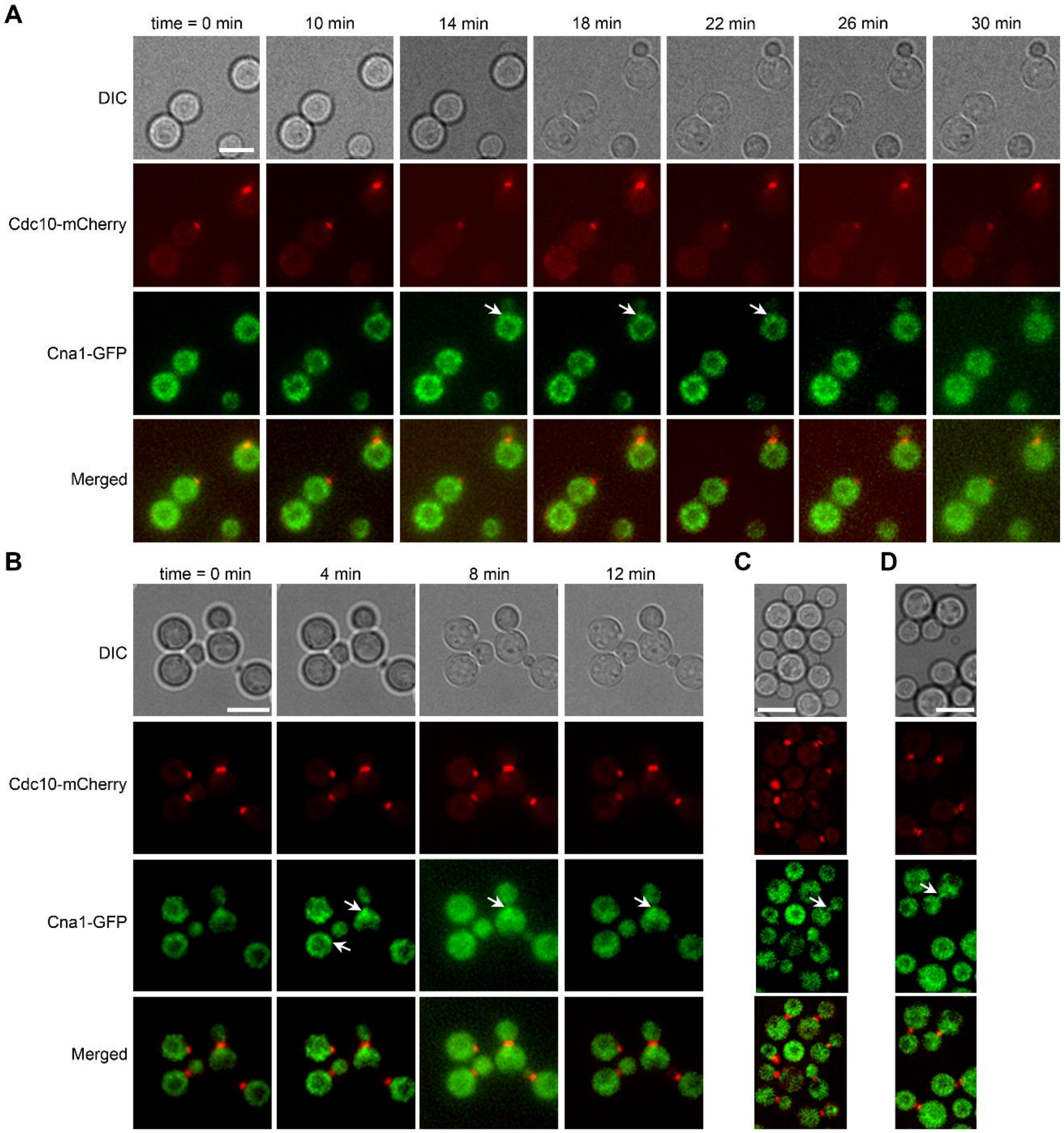


**Fig. S4. Cna1 co-localizes with Cdc10 at the mother-bud neck**. (A-B) Real-time live cell imaging showing the localization dynamics of Cna1-GFP in cells expression Cdc10-mCherry as a marker for mother bud neck. In budding cells, Cna1-GFP transiently localizes near the mother-bud neck and partially co-localizes with Cdc10 (marked with white arrows). Cna1-GFP also exhibited dynamic puncta localization across the cytoplasm that resembles calcineurin's previously reported localization at P-bodies/stress granules in *C. neoformans*. Scale bars, 5 µm. (C-D) Snapshots of cells showing co-localization of Cna1-GFP and Cdc10-mCherry at mother bud neck in dividing cells. Scale bars, 5 µm.

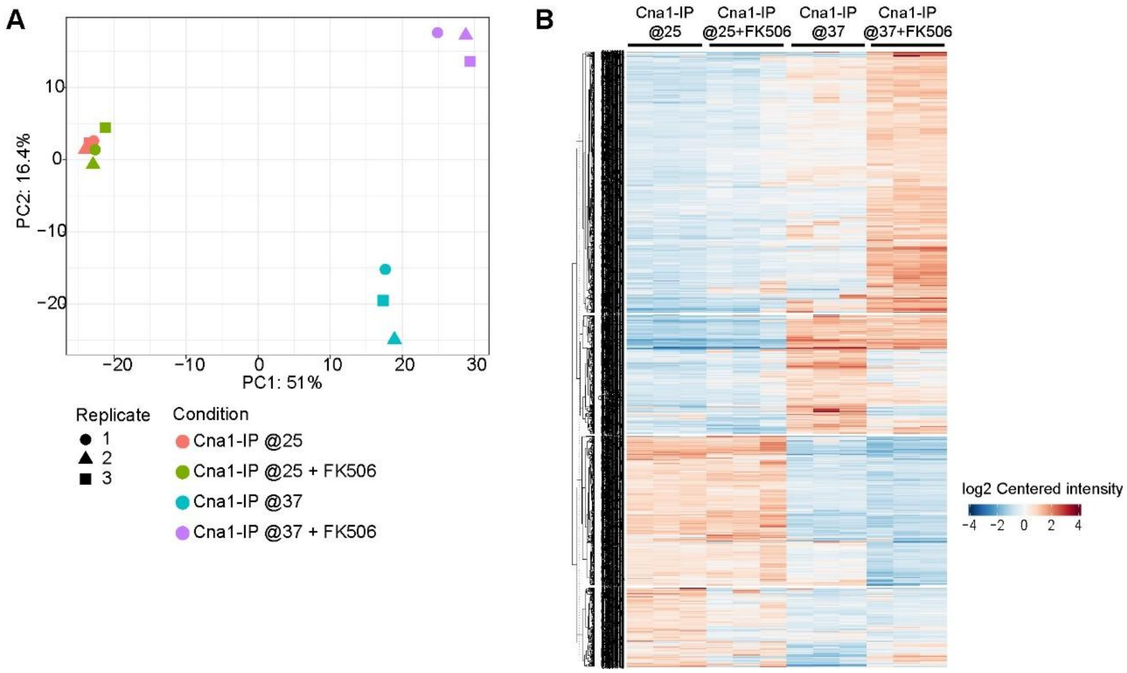


**Figure S5. Analysis of Cna1-GFP immunoprecipitation samples.** (A-B) A PCA plot and heatmap showing the sample correlation and replication of Cna1-GFP immuno-pulldown samples at different conditions.

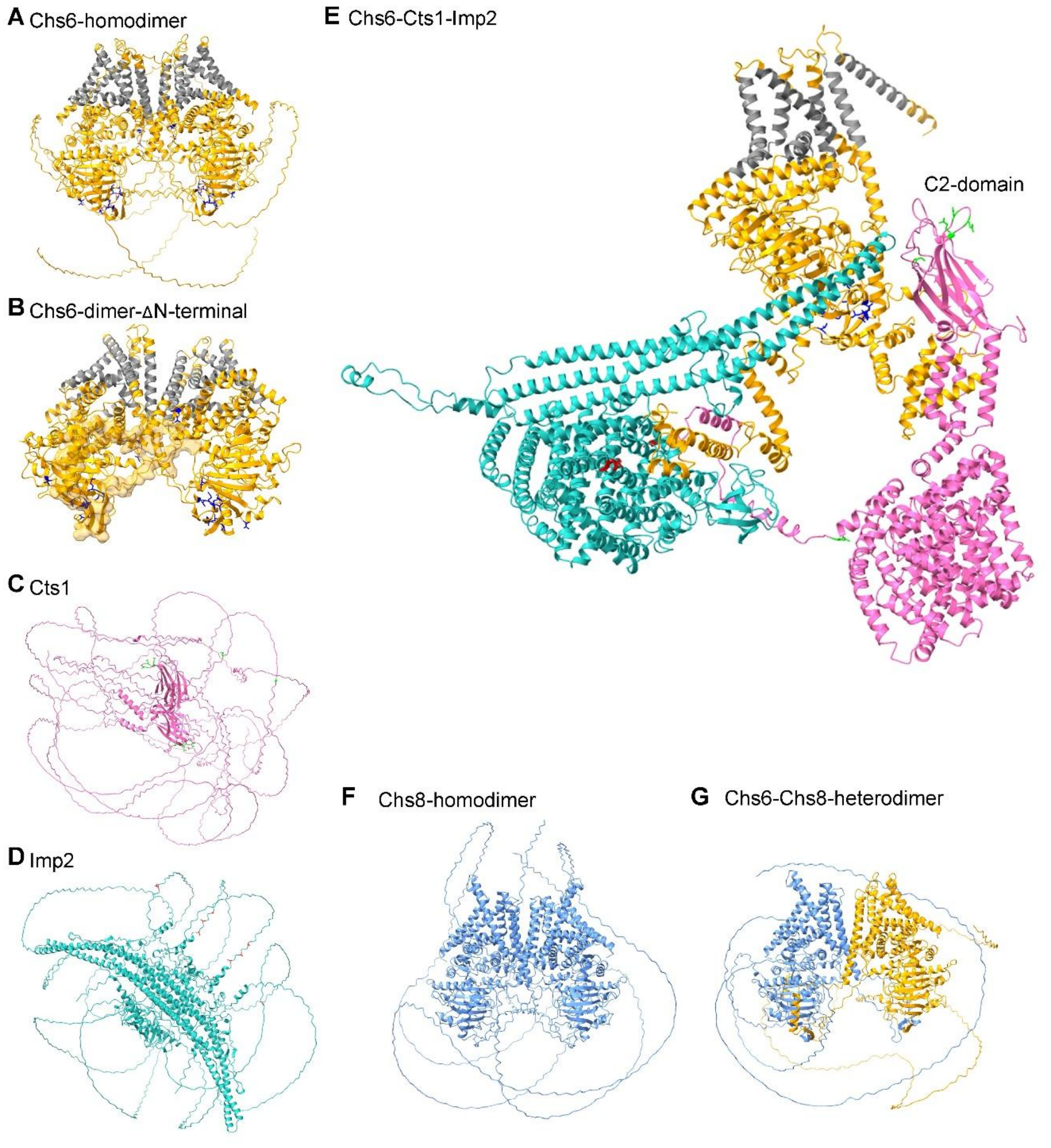


**Fig. S6. AlphaFold models for Chs6, Cts1, and Imp2.** (A) AlphaFold model for the Chs6 dimer predicted its transmembrane domains (highlighted in gray), and chitin synthase domain with the highly disordered domain N-terminal that remained unstructured. Identified calcineurin suppressor mutations are marked in blue. (B) Mapping of the domain swap region (highlighted as the surface cloud in one unit of the dimer) on the predicted Chs6 revealed that all identified mutations are either part of the domain swap region or present in close proximity to the domain swap region. The disordered N-terminal region is not shown. (C) AlphaFold prediction for Cts1 only predicted its C2-domain with the rest of the protein remaining unstructured. Identified mutations are marked in green color. (D) Imp2 structure prediction revealed a structure for the F-BAR domain with most of the protein being unstructured. All the identified mutations are present in the unstructured region and are marked with red color. (E) Multimer prediction of one unit of Chs6, Cts1, and Imp2 showed the presence of three proteins in a complex that resulted in the complete structuring of disordered regions observed in the A, C, and E models. Respective mutations for each of the proteins are highlighted in the same colors as in A, C, and E. (F-G) AlphaFold models for Chs8 homodimer and Chs6-Chs8 heterodimer revealed structure similarities between the two proteins and formation of potential heterodimer among the two.

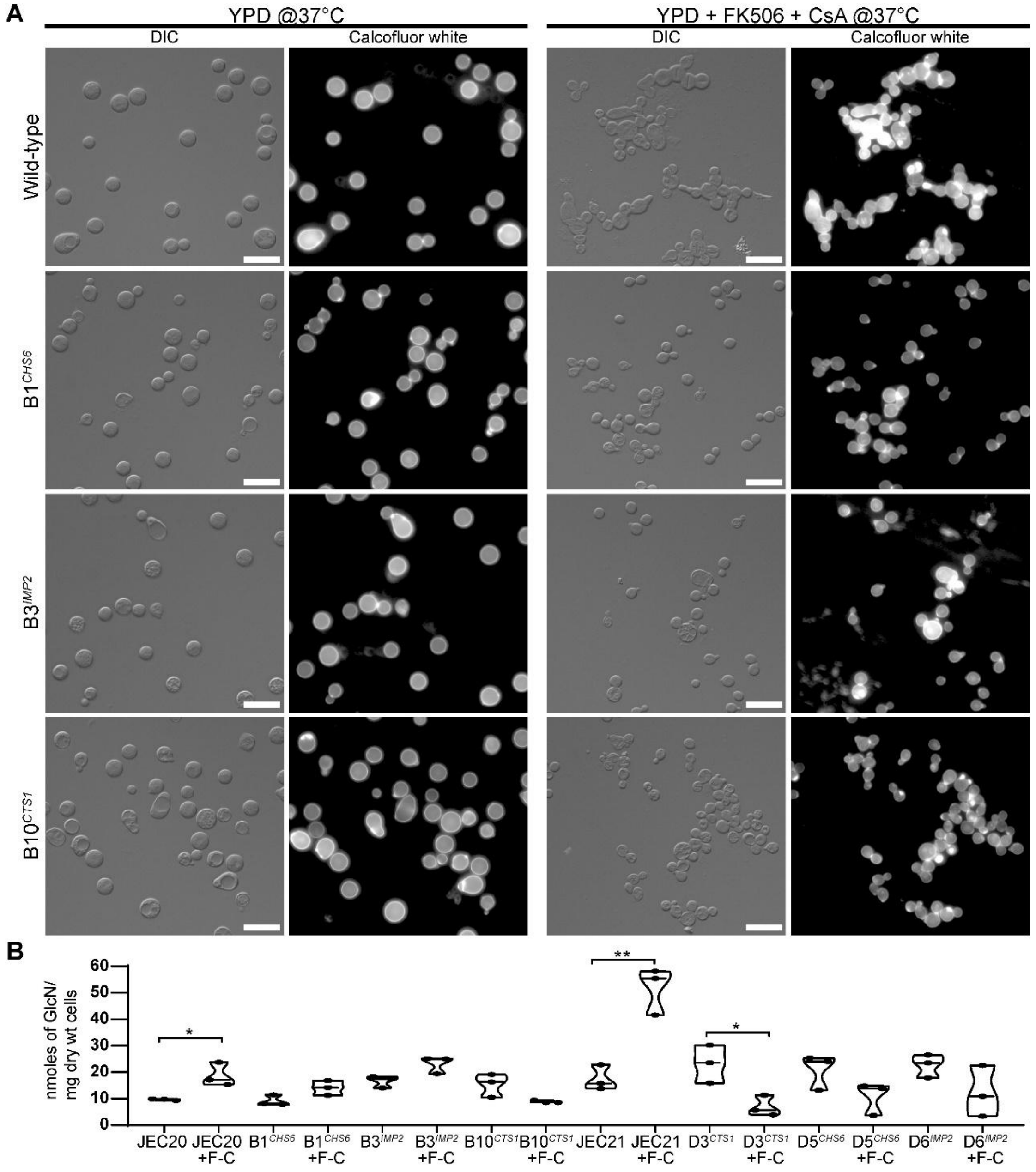


**Fig. S7. Calcineurin bypass mutations suppress budding defects and chitin accumulation phenotypes upon calcineurin inhibition.** (A) Images showing wild-type and calcineurin bypass mutant cells stained for chitin with calcofluor white both in the absence and presence of FK506 and CsA at 37°C. DIC images depict the presence of a large budded, chain of cells in the wild type that are restored to a normal state in the suppressor mutants. Scale bars, 20 µm. (B) Levels of chitin and chitosan were measured in the wild-type strains, JEC20 and JEC21, and their respective bypass mutants (B1, B3, B10 for JEC20 and D3, D5 and D6 for JEC21) with the MBTH assay. F-C refers to samples treated with FK506 and cyclosporin A. *, $p<0.05$; **, $p<0.01$; ns, non-significant.

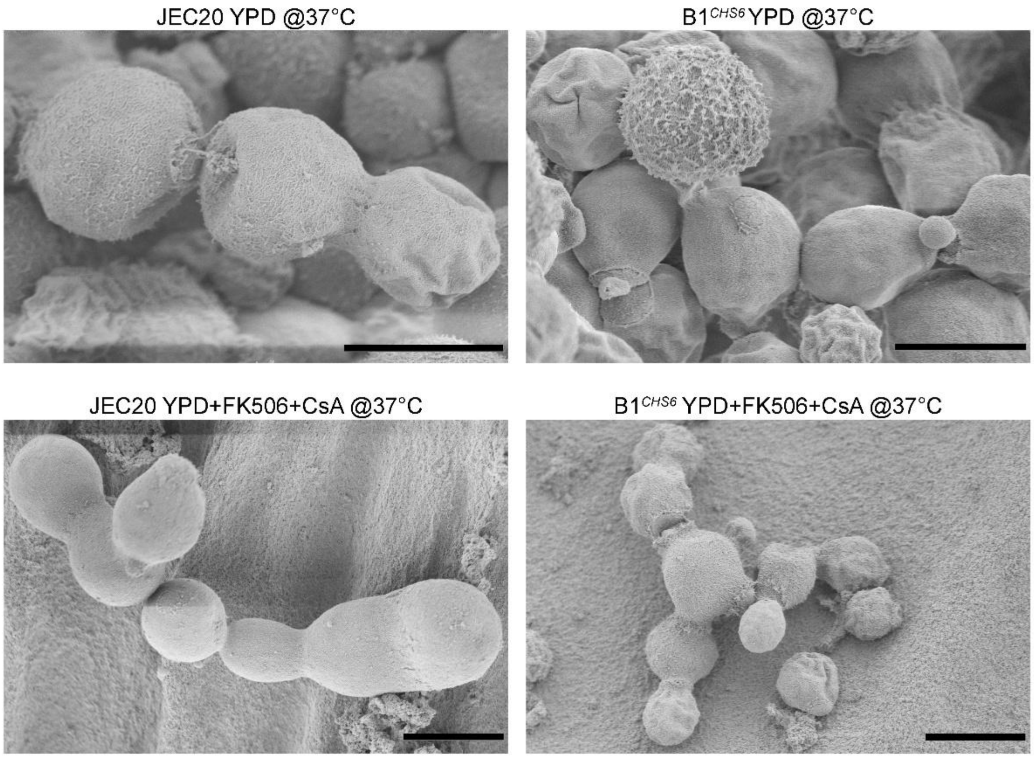


**Fig. S8. Scanning electron microscopy (SEM) revealed cytokinesis defect upon calcineurin inhibition.** SEM images showing cells at various stages in the wild-type JEC20 and its respective B1 suppressor with a mutation in *CHS6*. Upon calcineurin inhibition, the wild-type strain formed extensive chains of cells with broad necks revealing a failed cytokinesis event, which was largely restored in the suppressor mutant. Bars, 5 µm.

**Table S1. Summary of dominant/recessive phenotypes of selected suppressor diploids.**

| | Growth on YPD @37°C | Growth on YPD+Fk506 +CsA @37°C | Growth on SD-ura-ade | Growth on 5-FOA | Self-filamentation | Ploidy by Flow cytometry |
|---|---|---|---|---|---|---|
| **B1 (*CHS6*) x JEC50 diploids** | Yes | Yes | Yes | Yes | Yes | Diploid |
| | Yes | Yes | Yes | Yes | Yes | Diploid |
| | Yes | Yes | Yes | Yes | Yes | Diploid |
| | Yes | Yes | Yes | Yes | Yes | Diploid |
| | Yes | Yes | Yes | Yes | Yes | Diploid |
| | Yes | Yes | Yes | Yes | Yes | Diploid |
| | Yes | Yes | Yes | Yes | Yes | Diploid |
| | Yes | Yes | Yes | Yes | Yes | Diploid |
| **B3 (*IMP2*) x JEC50 diploids** | Yes | Yes | Yes | No | Yes | Diploid |
| | Yes | Yes | Yes | No | Yes | Diploid |
| | Yes | Yes | Yes | No | Yes | Diploid |
| | Yes | Yes | Yes | No | Yes | Diploid |
| | Yes | Yes | Yes | No | Yes | Diploid |
| | Yes | Yes | Yes | No | Yes | Diploid |
| | Yes | Yes | Yes | No | Yes | Diploid |
| | Yes | Yes | Yes | No | Yes | Diploid |
| **B10 (*CTS1*) x JEC50 diploids** | Yes | Yes | Yes | Yes | Yes | Diploid |
| | Yes | Yes | Yes | Yes | Yes | Diploid |
| | Yes | Yes | Yes | Yes | Yes | Haploid/Aneuploid |
| | Yes | Yes | Yes | Yes | Yes | Diploid |
| | Yes | Yes | Yes | Yes | Yes | Haploid/Aneuploid |
| | Yes | Yes | Yes | Yes | Yes | Diploid |
| | Yes | Yes | Yes | Yes | Yes | Diploid |
| | Yes | Yes | Yes | Yes | Yes | Diploid |
| **C12 (*ISC1*) x JEC171 diploids** | Yes | No | Yes | No | Yes | Diploid |
| | Yes | No | Yes | No | Yes | Diploid |
| | Yes | No | Yes | No | Yes | Diploid |
| | Yes | No | Yes | No | Yes | Diploid |
| | Yes | No | Yes | No | Yes | Diploid |
| | Yes | No | Yes | No | Yes | Diploid |
| | Yes | No | Yes | No | Yes | Diploid |
| | Yes | No | Yes | No | Yes | Diploid |
| **D6 (*IMP2*) x JEC171 diploids** | Yes | Yes | Yes | Yes | Yes | Diploid |
| | Yes | Yes | Yes | Yes | Yes | Diploid |
| | Yes | Yes | Yes | Yes | Yes | Diploid |
| | Yes | Yes | Yes | Yes | Yes | Diploid |
| | Yes | Yes | Yes | Yes | Yes | Diploid |
| | Yes | Yes | Yes | Yes | Yes | Diploid |

| | | | | | | |
|---|---|---|---|---|---|---|
| | Yes | Yes | Yes | No | Yes | Diploid |
| | Yes | Yes | Yes | Yes | Yes | Diploid |
| **F9 (*CHS6*) x JEC50 diploids** | Yes | No | Yes | Yes | Yes | Diploid |
| | Yes | No | Yes | No | No | Haploid/Aneuploid |
| | Yes | Yes | Yes | Yes | Yes | Diploid |
| | Yes | Yes | Yes | Yes | Yes | Diploid |
| | Yes | Yes | Yes | Yes | Yes | Diploid |
| | Yes | Yes | Yes | Yes | Yes | Diploid |
| | Yes | Yes | Yes | Yes | Yes | Diploid |
| | Yes | Yes | Yes | Yes | Yes | Diploid |
| **JEC20 x JEC50 diploids** | Yes | Yes | Yes | Yes | No | Haploid/Aneuploid |
| | Yes | No | Yes | Yes | Yes | Diploid |
| | Yes | Yes | Yes | Yes | No | Haploid/Aneuploid |
| | Yes | No | Yes | Yes | Yes | Diploid |
| | Yes | No | Yes | Yes | Yes | Diploid |
| | Yes | No | Yes | Yes | Yes | Diploid |
| | Yes | No | Yes | No | Yes | Diploid |
| | Yes | No | Yes | Yes | Yes | Diploid |

**Table S2. List of 66 proteins shared between calcineurin-mediated phosphoproteome and Cna1 interacting proteins identified by immunoprecipitation at 37°C.**

| Gene ID | Description |
|---|---|
| **CNA00570** | membrane protein, putative |
| **CNA02360** | expressed protein |
| **CNA02740** | putative nuclear polyadenylated mRNA-binding protein |
| **CNA02810** | RAS small monomeric GTPase, putative |
| **CNA03900** | eukaryotic initiation factor 4F subunit P130, putative |
| **CNA04800** | hypothetical protein |
| **CNA05020** | hypothetical protein |
| **CNA06440** | nucleoporin nsp1, putative |
| **CNA06790** | purine nucleotide biosynthesis-related protein, putative |
| **CNA07220** | imidazoleglycerol phosphate synthase, putative |
| **CNA07460** | hypothetical protein |
| **CNB01290** | Cyclophilin A |
| **CNB01520** | t-SNARE, putative |
| **CNB01970** | pre-mRNA splicing factor, putative |
| **CNB02230** | conserved hypothetical protein |
| **CNB03790** | heat shock protein, putative |
| **CNB04280** | 20S proteasome subunit, putative |
| **CNB05100** | hypothetical protein |
| **CNB05540** | proteasome subunit alpha type 1, putative |
| **CNC00160** | phosphopyruvate hydratase, putative |
| **CNC00390** | hypothetical protein |
| **CNC01700** | fumarate hydratase, putative |
| **CNC02040** | DNAj protein, putative |
| **CNC02320** | heat shock protein 70, putative |
| **CNC02470** | phosphatase, putative |
| **CNC03540** | cytoplasm protein, putative |
| **CNC04180** | dihydroxyacetone kinase 1, putative |
| **CNC04510** | conserved hypothetical protein |
| **CNC04930** | cytoplasm protein, putative |
| **CNC06560** | conserved hypothetical protein |
| **CND00130** | tRNA (cytosine-5-)-methyltransferase, putative |
| **CND02340** | allantoicase, putative |
| **CND02450** | dihydrolipoyllysine-residue acetyltransferase, putative |
| **CND03740** | aldo-keto reductase, putative |
| **CND05160** | heat shock protein HSP60, putative |

| | |
|---|---|
| **CNE00550** | conserved hypothetical protein |
| **CNE01830** | 4-aminobutyrate aminotransferase, putative |
| **CNE02610** | nucleoside-diphosphate kinase, putative |
| **CNE04560** | expressed protein |
| **CNF02100** | regulation of carbohydrate metabolism-related protein, putative |
| **CNF03260** | conserved hypothetical protein |
| **CNF03710** | methylenetetrahydrofolate dehydrogenase [NAD(+)] |
| **CNF04030** | hypothetical protein |
| **CNF04670** | conserved hypothetical protein |
| **CNF04720** | pim1 protein (poly(a)+ RNA transport protein 2), putative |
| **CNG00370** | expressed protein |
| **CNG00780** | cytoplasm protein, putative |
| **CNG03800** | conserved hypothetical protein |
| **CNG04210** | protein-methionine-S-oxide reductase, putative |
| **CNH01240** | cytochrome-b5 reductase, putative |
| **CNH02770** | hypothetical protein |
| **CNH03230** | arsenite-resistant protein ASR2, putative |
| **CNI02160** | hypothetical protein |
| **CNI02360** | NADPH dehydrogenase 2, putative |
| **CNI03350** | yeast yak1, putative |
| **CNJ00770** | hypothetical protein |
| **CNJ03153** | unspecified product |
| **CNK00280** | phosphoglycerate mutase, putative |
| **CNK00660** | RAB small monomeric GTPase, putative |
| **CNK01880** | C2H2 type zinc finger transcription factor |
| **CNK01890** | hypothetical protein |
| **CNL04280** | conserved hypothetical protein |
| **CNL05140** | conserved hypothetical protein |
| **CNL06310** | phosphatidylinositol transfer protein |
| **CNM02070** | heat shock protein, putative |
| **CNN00430** | phosphoglucomutase, putative |

**Table S3. List of strains used in this study.**

| Strain name | Description | Reference |
|---|---|---|
| **JEC21α** | Wild-type *MAT*α | (1) |
| **JEC20a** | Wild-type *MAT***a** | (1) |
| **JEC50** | JEC21α *ade2* | (2) |
| **JEC171** | JEC20a *lys2 ade2* | (3) |
| **ERB005** | H99α *cdc42Δ::NAT* | (4) |
| **VYD401-449 and VYD451-465** | Calcineurin suppressors (See details in Dataset S1) | This study |
| **VYD272** | 5-FOA-resistant derivative of suppressor B1 | This study |
| **VYD273** | 5-FOA-resistant derivative of suppressor B3 | This study |
| **VYD274** | 5-FOA-resistant derivative of suppressor B10 | This study |
| **VYD275** | 5-FOA-resistant derivative of suppressor C12 | This study |
| **VYD276** | 5-FOA-resistant derivative of suppressor D6 | This study |
| **VYD278** | 5-FOA-resistant derivative of suppressor F9 | This study |
| **VYD283** | 5-FOA-resistant derivative of suppressor JEC20 | This study |
| **VYD365** | JEC21α *CDC10-mCherry::NEO* | This study |
| **VYD369** | JEC20**a** *cdc42Δ::NEO* | This study |
| **VYD371** | JEC21α *cdc42Δ::NEO* | This study |
| **VYD373** | JEC21α *CDC10-mCherry::NEO CNA1-GFP::NAT* | This study |
| **VYD375** | JEC21α *cts1Δ::NEO* | This study |
| **VYD376** | JEC21α *chs6Δ::NEO* | This study |
| **VYD377** | JEC21α *imp2Δ::NEO* | This study |

| | | |
|---|---|---|
| **VYD378** | D3$^{CTS1}$ *cts1Δ::NEO* | This study |
| **VYD379** | D3$^{CTS1}$ *chs6Δ::NEO* | This study |
| **VYD380** | D3$^{CTS1}$ imp2Δ::*NEO* | This study |
| **VYD381** | D5$^{CHS6}$ *cts1Δ::NEO* | This study |
| **VYD382** | D5$^{CHS6}$ *chs6Δ::NEO* | This study |
| **VYD383** | D5$^{CHS6}$ *imp2Δ::NEO* | This study |
| **VYD384** | D6$^{IMP2}$ *cts1Δ::NEO* | This study |
| **VYD385** | D6$^{IMP2}$ *chs6Δ::NEO* | This study |
| **VYD386** | D6$^{IMP2}$ imp2Δ::*NEO* | This study |
| **VYD3122-3129** | B1 (*CHS6*) x JEC50 diploids | This study |
| **VYD3130-3137** | B3 (*IMP2*) x JEC50 diploids | This study |
| **VYD3138-3145** | B10 (*CTS1*) x JEC50 diploids | This study |
| **VYD3146-3153** | C12 (*ISC1*) x JEC171 diploids | This study |
| **VYD3154-3161** | D6 (*IMP2*) x JEC171 diploids | This study |
| **VYD3170-3177** | F9 (*CHS6*) x JEC50 diploids | This study |
| **VYD3194-3201** | JEC20 x JEC50 diploids | This study |

**Table S4. List of primers used in this study.**

| Name | Sequence | Purpose |
|---|---|---|
| JOHE51178 | CACATCTCAGATGCCATTTTACCA | *MAT* locus analysis |
| JOHE51179 | AGCTCTAAGTCATATGGGTTATAT | |
| JOHE51180 | CTTAATTCACAGCACCAGCCTA | |
| JOHE51181 | GGTCATCACAGTCAGTCACCAC | |
| JOHE52679 | CTCCATCTTCCAACAACC | *URA5* amplification and sequencing |
| JOHE52680 | CAAGCTTACAAACTTGTAATATG | |
| JOHE52681 | TTGGATCTTGTCGACAACGG | |
| JOHE52682 | TGATTAGCTAGACCTAAACCC | |
| JOHE53301 | GGAAATGTCGACTACTAAGC | *CHS6* ORF test PCR |
| JOHE53302 | CGTGAAGTGATGCTTACACG | |
| JOHE53303 | CAACGACGAGAAAGGAAACG | *CHS8* ORF test PCR |
| JOHE53304 | GATGTCTTCTACACCTTTTACC | |
| JOHE53305 | ACGAAACCTATTAAACGGTACG | *CTS1* ORF test PCR |
| JOHE53306 | CGTATCTTGGTATAGATGGAC | |
| JOHE53307 | GGACCTGAAAGCAAGCGC | *IMP2* ORF test PCR |
| JOHE53308 | CCTATACTGCTCCGCCATAG | |
| JOHE53309 | ATACGAATTTCTTCATCCTAGG | *ISC1* ORF test PCR |
| JOHE53310 | CAATAATTTTCCTCTCCATTAGC | |
| JOHE53383 | CATTCCTGATACTCGTATCCACTGC | Cdc10-mCherry tagging construct |
| JOHE53384 | AAGGATTCAAGCCCAGAAGAGGTCCG | |
| JOHE53385 | GACAATGTTAACAACGAAGGCTG | |
| JOHE53386 | CCTCGCCCTTGCTCACCATTGCCGTCGATGCAGATGCTTG | |
| JOHE53387 | CAAGCATCTGCATCGACGGCAATGGTGAGCAAGGGCGAGG | |
| JOHE53388 | CAGAAAGAATATTCATTCATCCAAGCTTGGTACCGAGCTC | |
| JOHE53389 | GAGCTCGGTACCAAGCTTGGATGAATGAATATTCTTTCTG | |
| JOHE53390 | CTCTCCTTCCTACTGCGGT | |
| JOHE53399 | CAAAGTGCGAAGAAGAGAAGGGCCTCTTCGCTATTACGCC | Cna1-GFP tagging construct |
| JOHE53400 | GGCGTAATAGCGAAGAGGCCCTTCTCTTCTTCGCACTTTG | |
| JOHE53401 | CCTCAACTGTTGCACCAAGGAG | |
| JOHE53402 | CTGCTAGGCCAAGGTAAGACTCC | |
| JOHE53403 | CTTTATGGATGTGTTCACCTGG | |
| JOHE53404 | CCTCGCCCTTGCTCACCATCTCTCTCTCGCCTTGACCGC | |
| JOHE53405 | GCGGTCAAGGCGAGAGAGAGATGGTGAGCAAGGGCGAGG | |
| JOHE53406 | GAGACTTAAGTCATGGAGGCTC | |
| JOHE53427 | ACCGGCAGGGTATACTGTTGAGGACGATATAGCACATAAGGTTTTA GAGCTAGAAATAGC | CdnCdc10-mCherry-gRNA-1 |
| JOHE53428 | ACCGGCAGGGTATACTGTTGACGTTAATAAGACAAAAGAGGTTTTA GAGCTAGAAATAGC | CdnCdc10-mCherry-gRNA-2 |

| | | |
|---|---|---|
| JOHE53431 | ACCGGCAGGGTATACTGTTGATTGTTGGGGCTACACAGTGGTTTTAGAGCTAGAAATAGC | CdCna1-GFP-gRNA-1 |
| JOHE53432 | ACCGGCAGGGTATACTGTTGAGAATTTAGGAAAGGGTCCAGTTTTAGAGCTAGAAATAGC | CdCna1-GFP-gRNA-2 |
| JOHE53782 | GTGCCCGCAGTCAAAGAGACGG | Cdc42 deletion construct |
| JOHE53783 | TGATTCACATAGCACAACGTCC | |
| JOHE53784 | gcggccgctctagaactagtgCAGTGACAGCAGCGCAGAAGAG | |
| JOHE53785 | CTCCAGCTCACATCCTCGCAGCCATGTCTGCATGGTTGGCTAGGAGG | |
| JOHE53786 | CCGTGTTAATACAGATAAACCGGCCTTGATCCAAGATACCCAACATTCC | |
| JOHE53787 | cgaattcctgcagcccgggAAGGGAGAATCTGTGTAAGATAGG | |
| JOHE53790 | ATGGCTGCGAGGATGTGAGCTGGAG | |
| JOHE53791 | GGCCGGTTTATCTGTATTAACACGG | |
| JOHE53788 | ACCGGCAGGGTATACTGTTGGTACCGGCGAGAGGCTCATGGTTTTAGAGCTAGAAATAGC | CdCdc42-del-gRNA1 |
| JOHE53789 | ACCGGCAGGGTATACTGTTGTATCTCGTAGGCCATCGTGGGTTTTAGAGCTAGAAATAGC | CdCdc42-del-gRNA2 |
| JOHE55540 | CCTTCAAACGTGATTCAGAGGC | Chs6 deletion construct |
| JOHE55541 | TATCGGTATCGGGCGTTCCAG | |
| JOHE55542 | AATGTGATCGTGAATGATAGCG | |
| JOHE55543 | CAGCTCACATCCTCGCAGCCATGATGTGGTGGATGTATGTATGG | |
| JOHE55544 | CCATACATACATCCACCACATCATGGCTGCGAGGATGTGAGCTG | |
| JOHE55545 | CTGATAACAAGGTACGCTGTTGAGCGGTTTATCTGTATTAACACGG | |
| JOHE55546 | CCGTGTTAATACAGATAAACCGCTCAACAGCGTACCTTGTTATCAG | |
| JOHE55547 | CCGCAAACTCAAGCACCAGC | |
| JOHE55548 | ACCGGCAGGGTATACTGTTGGAGGGAGGGTAGAAACTCGGGTTTTAGAGCTAGAAATAGC | Cdn-CHS6-del-gRNA-1 |
| JOHE55549 | ACCGGCAGGGTATACTGTTGCTTTCGTACCAATGTAAGCAGTTTTAGAGCTAGAAATAGC | Cdn-CHS6-del-gRNA-2 |
| JOHE55550 | AAGCTCCAGCAACTCATGGC | CTS1 deletion construct |
| JOHE55551 | AAGGTGTATTGCAGGACTGAGG | |
| JOHE55552 | CCGCAAGCTAGAAGCGAGAG | |
| JOHE55553 | CAGCTCACATCCTCGCAGCCATCCTTACGGGAGATGAGAAGAG | |
| JOHE55554 | CTCTTCTCATCTCCCGTAAGGATGGCTGCGAGGATGTGAGCTG | |
| JOHE55555 | GCTGAGGATGTAGCTGATTCACCGGTTTATCTGTATTAACACGG | |
| JOHE55556 | CCGTGTTAATACAGATAAACCGGTGAATCAGCTACATCCTCAGC | |
| JOHE55557 | GATCCGGAGATCATTGAGGAC | |
| JOHE55558 | ACCGGCAGGGTATACTGTTGGGAAGTGGTTGAGAATATTGGTTTTAGAGCTAGAAATAGC | Cdn-CTS1-del-gRNA-1 |
| JOHE55559 | ACCGGCAGGGTATACTGTTGTCATATCTCCAATATCACAGGTTTTAGAGCTAGAAATAGC | Cdn-CTS1-del-gRNA-2 |
| JOHE55560 | GCGATCACTGATATGACTGC | IMP2 deletion construct |
| JOHE55561 | CACTGAGTCATGCATTCCAGC | |
| JOHE55562 | GTCGACGAGAGGCAGAAACG | |

| | | |
|---|---|---|
| JOHE55563 | CAGCTCACATCCTCGCAGCCATAGCGTGACGTTGTTCTAAGC | |
| JOHE55564 | GCTTAGAACAACGTCACGCTATGGCTGCGAGGATGTGAGCTG | |
| JOHE55565 | CCATTCGATACAGCCGAAATGCGGTTTATCTGTATTAACACGG | |
| JOHE55566 | CCGTGTTAATACAGATAAACCGCATTTCGGCTGTATCGAATGG | |
| JOHE55567 | AAGCCAGACGTTATGAGAG | |
| JOHE55568 | ACCGGCAGGGTATACTGTTGAGCACTCACAGAACATGACAGTTTT AGAGCTAGAAATAGC | Cdn-IMP2-del-gRNA-1 |
| JOHE55569 | ACCGGCAGGGTATACTGTTGCTATACAATGGACCAAACGGGTTTTA GAGCTAGAAATAGC | Cdn-IMP2-del-gRNA-2 |

**Dataset S1 (separate file). List of all suppressor mutants describing the background identified genetic change and their impact on protein-coding.**

**Dataset S2 (separate file). List of genes present in all identified aneuploid regions.**

**Dataset S3 (separate file). List of phosphosites differentially enriched between 25°C and 37°C for both the wild-type and calcineurin mutant strains Type or paste legend here.**

### SI References


1. J. Heitman, B. Allen, J. A. Alspaugh, K. J. Kwon-Chung, On the origins of congenic *MATα* and *MAT***a** strains of the pathogenic yeast *Cryptococcus neoformans*. *Fungal Genet Biol* **28**, 1-5 (1999).
2. C. B. Nichols, J. A. Fraser, J. Heitman, PAK kinases Ste20 and Pak1 govern cell polarity at different stages of mating in *Cryptococcus neoformans*. *Mol Biol Cell* **15**, 4476-4489 (2004).
3. K. B. Lengeler, G. M. Cox, J. Heitman, Serotype AD strains of *Cryptococcus neoformans* are diploid or aneuploid and are heterozygous at the mating-type locus. *Infect Immun* **69**, 115-122 (2001).
4. E. R. Ballou, C. B. Nichols, K. J. Miglia, L. Kozubowski, J. A. Alspaugh, Two CDC42 paralogues modulate *Cryptococcus neoformans* thermotolerance and morphogenesis under host physiological conditions. *Mol Microbiol* **75**, 763-780 (2010).
